# NoroScope: Exploring the Mutational Landscape of the Human Norovirus Capsid Protein with Context-Aware Machine Learning

**DOI:** 10.64898/2026.09.17.752373

**Authors:** Sebastian Bowyer, David J. Allen, Nicholas Furnham

## Abstract

Anticipating the effects of mutations in viral proteins is important for the development of therapeutics and control strategies but remains challenging as mutation phenotypes are shaped by complex overlapping sequence, structural and evolutionary constraints. Machine-learning approaches to mutation-effect prediction have shown considerable success for viruses with exceptionally rich training datasets, but their application to the majority of viruses with comparatively limited data remains underexplored. Here we introduce NoroScope, an interpretable machine-learning framework designed to prioritise plausible amino-acid substitutions in the human norovirus GII.4 capsid protein VP1 through integration of complementary biological evidence. Using VP1 protein sequences and experimentally resolved structures as a foundation, we generated datasets comprising zero-shot mutation scores from the Evolutionary Scale Modeling 2 protein large language model, *in silico* deep-mutational-scanning estimates of mutation-associated effects on protein stability and histo-blood group antigen binding, and descriptors of the evolutionary history of VP1. NoroScope integrated these evidence sources to prioritise historically observed alternative amino acids according to whether they subsequently received recurrent support in natural virus populations. Across retrospective 2011 and 2016 prediction tasks, the complete model achieved average precision values of 0.805 and 0.834 and recovered 87.9% and 92.9% of future-supported candidates within the top five alternatives at their respective positions. Feature ablation showed that evolutionary, sequence-model and structural information provided distinct and complementary predictive signals, with their integration supporting accurate amino acid prioritisation and revealing biologically informative disagreements between evidence sources. Experimental analysis at VP1 position 297 further supported molecular predictions for selected substitutions in a virus-like particle assembly system. Together, these results establish NoroScope as a framework for integrating pretrained protein representations with target-specific molecular and evolutionary context to explore viral mutation landscapes where available data are comparatively limited.

## Introduction

Viruses and particularly RNA viruses are capable of rapid evolution through genetic variation. Amino-acid substitutions in viral proteins are able to alter phenotypes including antigenicity, host interactions, transmissibility, antiviral sensitivity and immune escape (1,2). Predicting the consequences of individual substitutions is challenging as their effects are governed by multiple overlapping constraints. At the molecular level, substitutions must remain compatible with core properties including protein stability, structure, molecular interactions and the surrounding genetic background, while persistence at the population level is shaped by host immunity, transmission, natural selection, genetic drift and epidemiological opportunity (2,3). Recent advances in machine learning (ML) techniques provide increasingly powerful tools for investigating such relationships, including approaches capable of learning from high-dimensional biological data, integrating heterogeneous sources of information and transferring representations learned from large biological datasets to new prediction tasks (4–6).

The potential of these approaches has been demonstrated most thoroughly with SARS-CoV-2, as a uniquely large combination of genomic, experimental and structural data became available as the result of a coordinated global effort during the COVID-19 pandemic. EVEscape, for example, integrates evolutionary sequence modelling with structural and biophysical information to prioritise mutations with potential for immune escape, while CoVFit combines a protein language model with extensive coronavirus-specific sequence and phenotype information to model SARS-CoV-2 fitness (7,8). However, the exceptional depth of sequence and experimental information available for SARS-CoV-2 is not representative of the state of research for most studied viruses. This raises the question of how emerging computational approaches can be applied to viruses with comparatively limited data availability. To investigate this path, we selected human norovirus (HuNoV) as a representative virus.

HuNoVs are positive-sense, single-stranded RNA viruses and a leading cause of acute gastroenteritis worldwide, responsible for substantial medical and economic burden as well as frequent outbreaks in both community and institutional settings (9). Within HuNoV, the GII.4 genotype has historically accounted for a majority of global outbreaks and has evolved within the global human population through the emergence of successive, antigenically distinct epidemic variants (10,11). VP1, the major capsid protein of HuNoV, is central to this evolutionary history. VP1 forms the viral capsid and mediates several important interactions within the host. The protein consists of a shell (S) domain and surface-exposed protruding (P) domain, with the latter containing the hypervariable P2 subdomain in which major epitopes and the histo-blood group antigen (HBGA) binding interface are located (12,13). Amino-acid variation within VP1 and particularly within P2 has been repeatedly associated with altered antigenic recognition, changes in HBGA interactions and the diversification and emergence of GII.4 variants (11,14,15).

In this work we introduce NoroScope: an interpretable, context-aware framework for studying the molecular and evolutionary interactions that shape amino-acid substitutions in viral proteins. NoroScope uses ML techniques to integrate complementary sources of biological information into a ranking tool which prioritises plausible alternative amino acids at each position within a target protein. Using a dataset of aligned protein sequences as a base, the initial molecular input to the framework is provided by Evolutionary Scale Modeling 2 (ESM-2), a foundation protein large language model (pLLM) which generates sequence-dependent scores describing the relative plausibility of alternative amino acids (6). To supplement sequence-based representations provided by ESM-2, NoroScope integrates target-specific information that describes complementary constraints on GII.4 VP1 evolution. These data include computational estimates of mutation-associated effects on protein stability as well as the binding energetics of the VP1-HBGA complex alongside measures of historical amino-acid variation and recurrence derived from time-aware sequence data and annotations of established functional regions. The resulting features are integrated using supervised ML to generate relative rankings of alternative amino acids within individual VP1 positions. Here, we evaluate NoroScope retrospectively across historical stages of GII.4 evolution, testing whether relationships learned between molecular and evolutionary context can generalise to protein positions withheld from model training and prioritise substitutions that subsequently receive recurrent support in natural virus populations. Through feature ablation, control analyses and targeted experimental validation, we additionally examine how these different evidence sources contribute to model behaviour and how interpretable disagreements between them can unlock key biological insights.

## Results

### A longitudinal sequence dataset captures GII.4 VP1 evolution

The NoroScope modelling framework requires sequences with known sample collection dates in order to connect historical sequence context to later mutation support. The foundation for this dataset comes from the NOROPATROL project, which collated and sequenced full HuNoV genomes from patient samples collected in England and Wales between the years of 1994 and 2023 (16). The NoroScope sequence dataset additionally integrates complete GII.4 VP1 sequences obtained from NCBI Virus (17).

Following multiple sequence alignment, the common NoroScope coordinate system was established to ensure that all VP1 protein sequences were the correct length of 540 or 539 amino acids with one canonical indel permitted – that being the single amino acid insertion at position 394 which emerged with the Farmington Hills variant in 2002 (18). The final NoroScope modelling dataset contained 1,657 unique, dated entries after final quality control, of which 1,009 originated from NCBI Virus and 648 originated from NOROPATROL.

We identified three distinct functional groups for annotation: epitopes, the HBGA-binding pocket and the “breathing core”. Residues associated with these functional categories were collated from published structural, antigenic and population-genomic studies (Table 1).

**Table 1:** Functional-region annotations used for HuNoV GII.4 VP1 in the NoroScope study. Residue sets were compiled from published structural, antigenic and population-genomic studies and mapped to the common NoroScope VP1 coordinate system, which reflects canonical protein numbering. Where published definitions differed between studies, the annotation retained the combined set of reported residues. Descriptions summarise the established or proposed structural and functional roles of each region.

| Name | Residues | Description | Reference |
| --- | --- | --- | --- |
| **Epitope A** | 294-299, 368, 372-373 | A highly exposed antigenic site on the surface of the P2 subdomain. Variation within this region has been associated with escape from blockade-antibody recognition and the emergence of new GII.4 variants. | Lindesmith *et al.*, 2012; Ford-Siltz *et al.*, 2022; Zheng *et al.*, 2022 (13,19,20) |
| **Epitope B** | 333, 382, 389 | A largely buried antigenic region located near the interface between VP1 monomers. Variation within this site has been proposed to influence the conformation or exposure of more surface-accessible epitopes. | Lindesmith *et al.*, 2012; van Loben Sels and Green, 2019; Ottosson *et al.*, 2022 (13,21,22) |
| **Epitope C** | 339-341, 375-378 | Located on the lateral surface of the P2 subdomain, adjacent to the HBGA-binding pocket. Variation within this region has been associated with the diversification and replacement of GII.4 variants. | Tohma *et al.*, 2019 (11) |
| **Epitope D** | 391-397 | A surface-exposed antigenic region situated along the ridge of the HBGA-binding pocket. Residues within this site contribute to both blockade-antibody recognition and modulation of HBGA-binding interactions. | Lindesmith *et al.*, 2022 (16) |
| **Epitope E** | 407, 411-414 | Located lateral to epitopes A and D on the P2 subdomain and less prominently exposed on the outer capsid surface. Variation within this region has been associated with GII.4 antigenic change. | Tohma *et al.*, 2019; Ford-Siltz *et al.*, 2022 (11,19) |
| **Epitope F** | 327, 404 | A conserved conformational blockade epitope. Antibody access to this site is regulated by particle conformation and by distal residues within the NERK motif. | Lindesmith *et al.*, 2018; Mallory *et al.*, 2019 (23,24) |
| **Epitope G** | 352, 355-357, 359, 364 | A surface-exposed antigenic region identified through population-genomic analysis. Variation within this site has been associated with the emergence and circulation of GII.4 variants. | Tohma *et al.*, 2019 (11) |
| **Epitope H** | 309, 310 | A putative surface-exposed antigenic motif identified through population-sequence analysis. Direct experimental mapping of antibody recognition at this site remains limited. | Tohma *et al.*, 2019 (11) |
| **Epitope I** | 402-403, 504, 506 | A structurally conserved conformational epitope recognised by broadly reactive GII.4-blocking antibodies. | Lindesmith *et al.*, 2019 (25) |
| **Breathing core (NERK motif and residue 234)** | 234, 310, 316, 484, 493 | Comprises the conserved NERK motif at residues 310, 316, 484 and 493 together with residue 234. These spatially separated residues regulate antibody access to occluded epitopes through effects on particle conformation. | Lindesmith *et al.*, 2018 (23) |
| **HBGA binding pocket: Loop 1** | 372-378 | A surface-exposed glycan-binding region located on the P2 subdomain of VP1. The pocket comprises three loops and an adjacent β-sheet that together form the structural interface for HBGA recognition. The β-sheet contributes to the comparatively conserved core of the binding site, while variation within the surrounding loops can alter glycan-binding specificity and affinity. | Shanker *et al.*, 2011, Liang *et al.*, 2021 (12,15) |
| **HBGA binding pocket: Loop 2** | 390-396 |  |  |
| **HBGA binding pocket: Loop 3** | 441-445 |  |  |
| **HBGA binding pocket: β-sheet** | 341-346 |  |  |

We next characterised amino acid variability across VP1 using positional Shannon entropy. Broadly, entropy patterns were consistent with established understanding of VP1 structure: the majority of positions were highly conserved, with greater variability concentrated within the P domain and particularly the P2 subdomain (Fig. 1). Variable, high-entropy positions were found within surface-exposed epitopes and the HBGA-binding pocket, consistent with the concentration of evolutionary change within exposed P2 loops.

**Figure 1:**
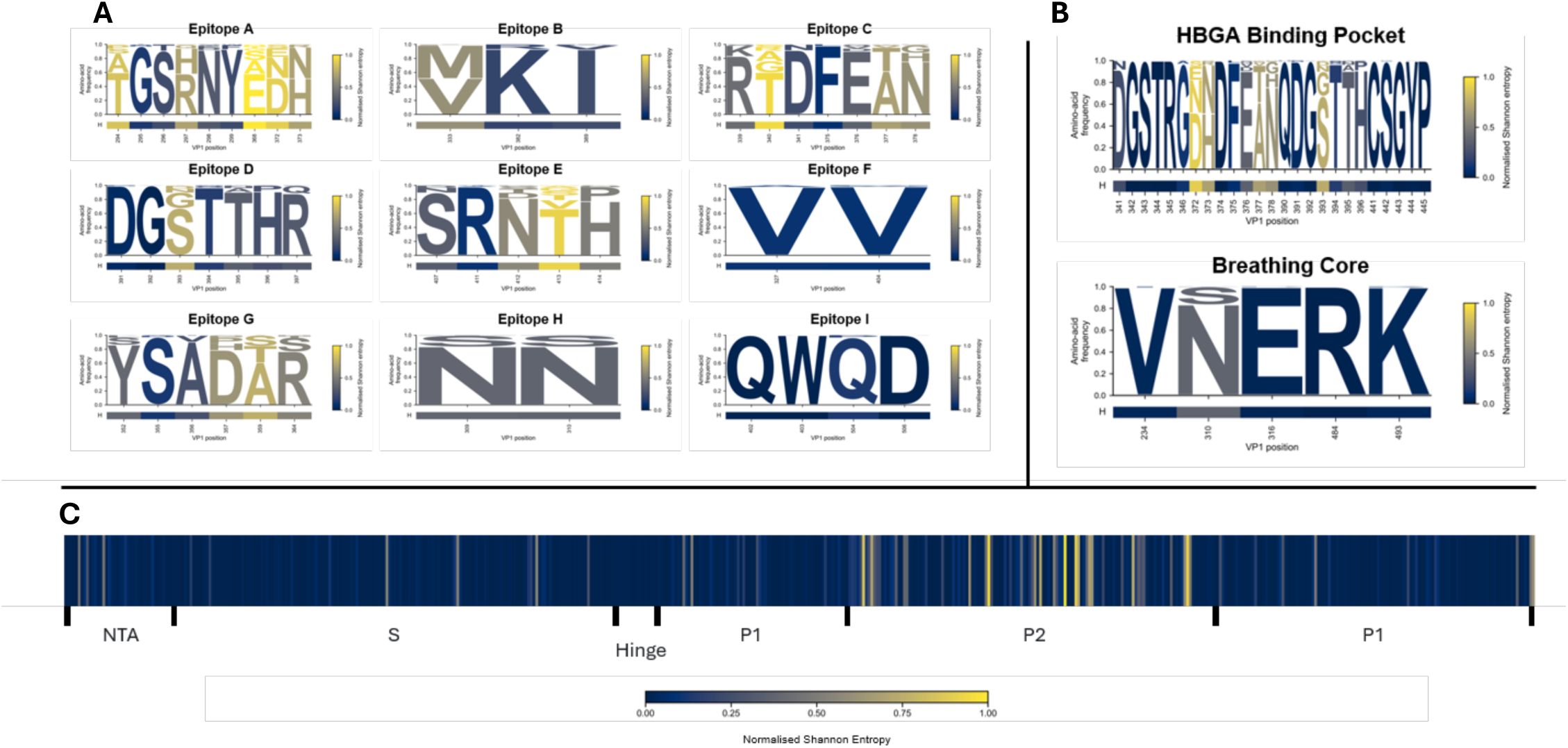
Sequence logo plots and full-protein entropy map of the NoroScope sequence dataset. Normalised Shannon entropy is displayed as a heatmap, with blue indicating low entropy and yellow indicating high entropy. **A:** Sequence logo plots displaying amino acid frequency and entropy for each residue within the nine epitopes annotated in the NoroScope dataset. Larger letters indicate a higher within-position frequency for the corresponding residue. **B:** Sequence logo plots for the HBGA binding pocket as well as the breathing core. **C:** A full-protein entropy map of the NoroScope sequence dataset. Per-position entropy is displayed as a heatmap across all 540 positions in the GII.4 VP1 protein sequence. Major structural domains are also labelled; the N-terminal arm (NTA), S domain, and P domain with P1 and P2 subdomains. A flexible hinge separates the S and P domains.

### Molecular and evolutionary datasets describe distinct constraints

#### Zero-shot prediction with a protein large language model captures biophysical plausibility

ESM-2, a family of pLLMs developed by Meta AI, was selected for the NoroScope zero-shot prediction task (6). Before expanding the model task across the complete dataset, we compared four architecture sizes using a representative GII.4 VP1 sequence and a broader set of variant medoids.

Quantitative analysis found that the 15B model achieved the strongest recovery of observed amino acids and the highest correspondence between model-predicted and empirical positional entropy across GII.4 variants (Fig. S1). The 15B model was taken forward for generation of log-likelihood ratios (LLRs) across the complete NoroScope sequence dataset.

Applying the 15-billion-parameter ESM-2 model across 1,657 sequences, 540 VP1 positions and 20 candidate amino acids generated 17,894,580 individual LLRs after accounting for the canonical 394 insertion, with each score representing one candidate amino acid at a given position within a specific sequence context. Within this framework, the same substitution at the same position could receive a different score in an alternative VP1 sequence background.

#### Preparation of GII.4 VP1 protein structures for in silico deep mutational scanning

*In silico* deep mutational scanning (DMS) was used to construct two complementary datasets describing predicted mutation-associated effects on the GII.4 VP1 P-domain dimer: one estimating changes in protein stability and the other estimating changes in the binding energetics of the dimer in complex with an HBGA type A glycan. Combining three experimental structures with four virus-like particle (VLP) sequence backgrounds generated 12 intended structure–sequence inputs for each DMS task.

#### In silico DMS identifies position-specific stability constraints in the GII.4 VP1 P-domain

The MutateX pipeline was used to automate saturation mutagenesis and ΔΔG calculation for each structure–sequence input in a protein stability scan of the GII.4 VP1 P domain (26). The scan generated 67,280 ΔΔG calculations across 11 structure–sequence inputs after one input was excluded as a technical outlier (Fig. S2).

Most candidate substitutions were predicted to be destabilising. Continuous stability estimates were highly consistent across experimental structures and VLP sequence backgrounds, with median pairwise Spearman correlations ranging from 0.927 to 0.971 (Fig. 2). Stability classifications were unanimous across all four VLP sequence backgrounds for 78.7% of candidate substitutions and majority-supported for a further 14.2%, indicating that the broad FoldX-derived stability landscape was robust to both structural-template and sequence context.

**Figure 2:**
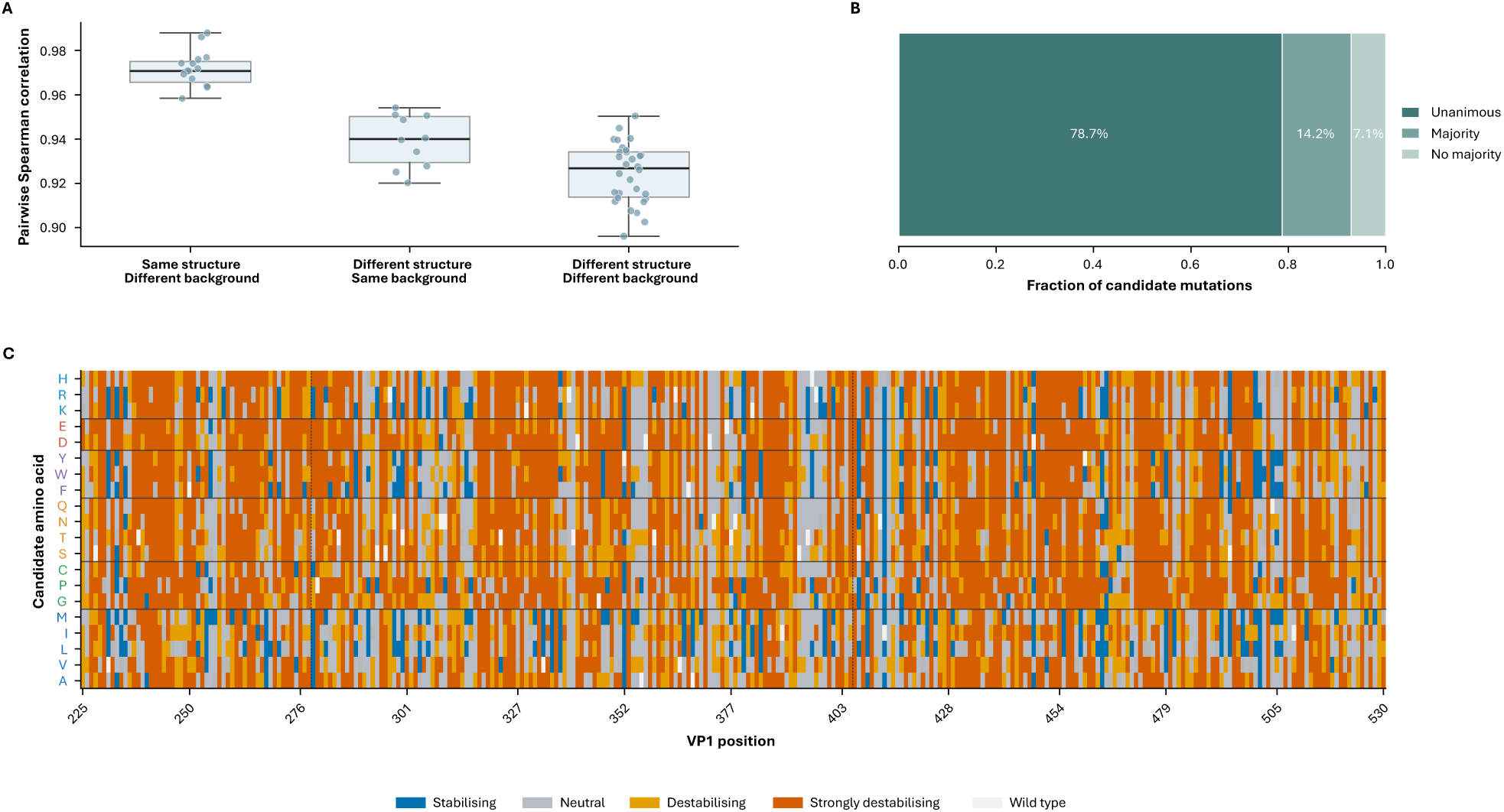
FoldX-derived stability landscape and robustness across experimental structures and VLP sequence backgrounds. A: Effects of experimental template and back-mutated sequence background on continuous FoldX ΔΔG estimates. Pairwise Spearman correlations are shown for DMS inputs sharing the same experimental template, sharing the same back-mutated VLP sequence background, or differing in both. Points represent individual pairs of structure–sequence inputs. **B:** Robustness of FoldX stability classifications across VLP sequence backgrounds. Candidate substitutions were classified as unanimous when all available backgrounds received the same stability category, majority-supported when more than half agreed, or lacking a majority otherwise. **C:** Majority stability category assigned to each candidate amino-acid substitution across the four VLP sequence backgrounds at structurally represented positions within the GII.4 VP1 P domain. Amino acids are grouped by biochemical similarity. White cells indicate the wild-type amino acid. Dashed vertical lines delimit the P2 subdomain.

#### In silico DMS identifies position-specific HBGA binding constraints

To estimate mutation-associated effects on the VP1–HBGA interaction, a paired wild-type–mutant workflow was established using Rosetta (27). Most candidate substitutions were predicted to have limited effects on the locally constrained VP1–HBGA complex, with 74.9% classified as binding-neutral. Predicted effects were strongly position-dependent, with most non-neutral classifications concentrated within the HBGA-binding pocket or the structural core of the P-domain dimer.

Continuous Rosetta estimates were more dependent on structural-template and sequence context than the FoldX stability estimates. Median pairwise Spearman correlations were 0.373 between calculations sharing the same experimental template, 0.338 between those sharing the same VLP sequence background and 0.301 when both contexts differed (Fig. 3). Despite this variation in the precise energetic ranking of substitutions, the direction of effect agreed in approximately 88% of pairwise comparisons. Binding-effect classifications were unanimous across sequence backgrounds for 73.0% of candidate substitutions and majority-supported for a further 16.3%.

**Figure 3:**
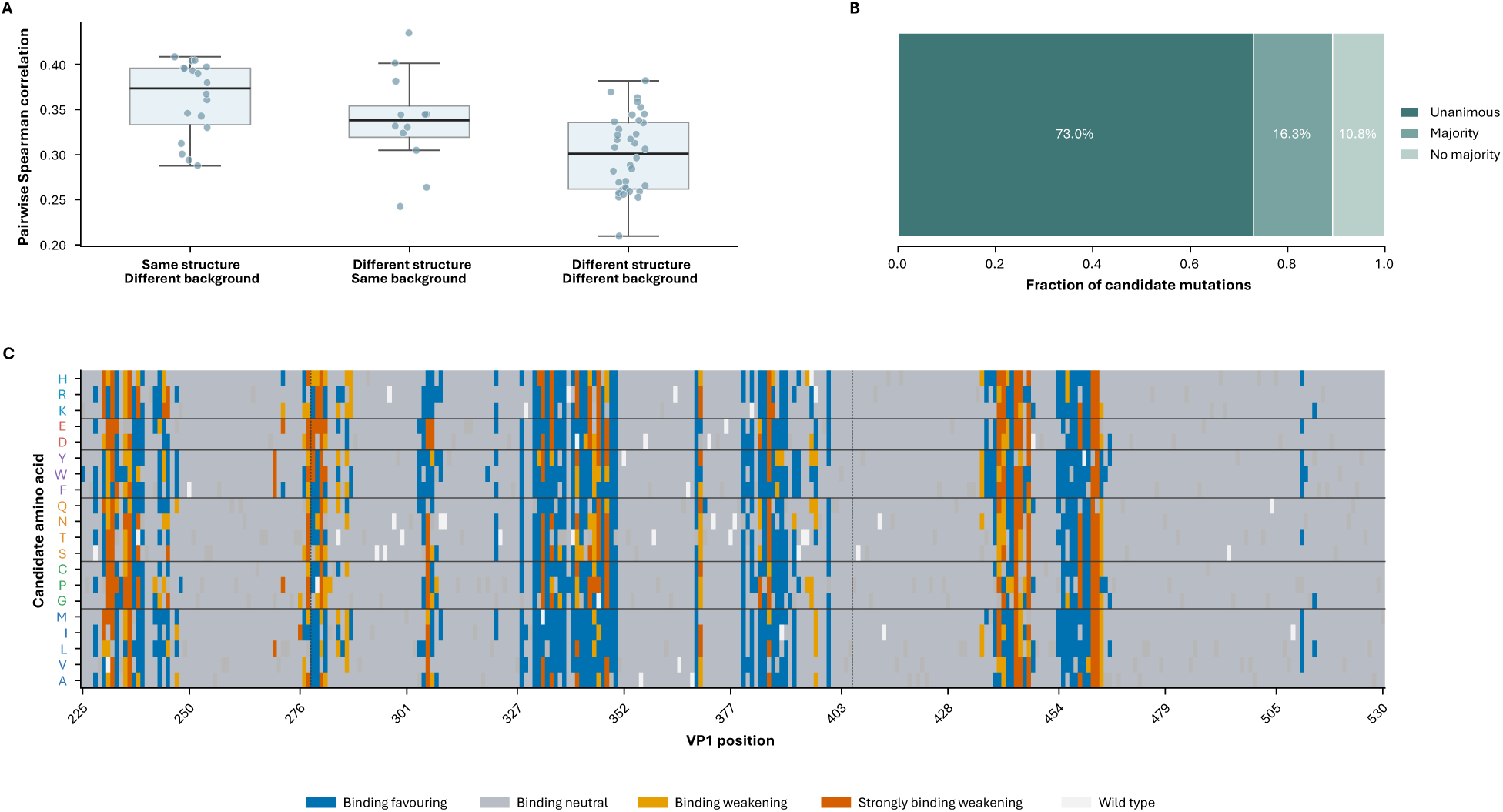
Rosetta-derived HBGA-binding landscape and robustness across experimental structures and VLP sequence backgrounds. A: Effects of experimental template and back-mutated sequence background on continuous Rosetta estimates. Pairwise Spearman correlations are shown for DMS inputs sharing the same experimental template, sharing the same back-mutated VLP sequence background, or differing in both. Points represent individual pairs of structure–sequence inputs. **B:** Robustness of Rosetta binding-effect classifications across VLP sequence backgrounds. Consensus effects were classified as non-neutral when the magnitude of ΔΔG_bind_ exceeded both 0.25 Rosetta energy units (REU) and twice the median replicate standard error of the mean (SEM); supported effects were classified as binding-favouring (<−0.25 REU), binding-weakening (>+0.25 to <+2 REU), or strongly binding-weakening (≥+2 REU), with remaining eBects classified as binding-neutral. Candidate substitutions were classified as unanimous when all available backgrounds received the same binding-effect class, majority-supported when more than half agreed, or lacking a majority otherwise. **C:** Majority binding-effect class assigned to each candidate amino-acid substitution across the four VLP sequence backgrounds at structurally represented positions within the GII.4 VP1 P domain. Amino acids are grouped by biochemical similarity. White cells indicate the wild-type amino acid. Dashed vertical lines delimit the P2 subdomain.

#### Evolutionary history captures recurrent and temporally structured mutation patterns

Although candidate amino acids can initially be divided according to whether they had been observed before a given cutoff, a simple binary “seen/unseen” label does not adequately describe their historical status. The Evolutionary Context Dataset (ECD) comprises features describing the historical distribution of candidate amino acids at each VP1 position before a defined temporal cutoff. Rather than recording only whether a candidate had previously been observed, the feature set summarises amino-acid frequency, temporal breadth, age and recency. ECD features were divided into site-level context and direct candidate-recurrence history. Site-level context described the historical variability and most common amino acid at each position, whereas direct candidate recurrence described the frequency and temporal distribution of each previously observed amino acid at that position.

ECD features were incorporated into NoroScope with all values calculated using only sequences sampled on or before the relevant historical cutoff.

### Integrating heterogeneous and evolutionary evidence into a temporally-aware mutation database

Each feature included within the NoroScope database was aligned to a common 540-position VP1 coordinate system and integrated around the unit of historical cutoff, VP1 position and candidate amino acid. This structure allowed the same candidate residue to acquire different feature values at alternative historical cutoffs as the available sequence contexts and recurrence history changed.

2011 and 2016 were selected as retrospective historical cutoff years for model predictions. For each cutoff, only sequences sampled on or before the selected year were permitted to contribute predictive sequence contexts, molecular feature aggregates or evolutionary-history features. This ensured that future mutation observations could not enter the predictive feature set.

#### NoroScope is a position-held-out mutation ranking model

The NoroScope model was formulated as a retrospective candidate mutation-ranking task. For a given historical cutoff year, the model assigns a relative score to alternative amino-acid states at each position in VP1 using the molecular, structural and evolutionary features available at that cutoff. At each position, the amino acid most frequently observed at the cutoff, termed the dominant amino acid, is excluded from the prediction task. The remaining 19 amino acids form the complete alternative-candidate pool.

The primary modelling cohort is then further restricted to non-dominant candidates that had been observed at least once before the cutoff (fig. S4). The primary model outcome was recurrent natural support during the five years following each historical cutoff. A candidate was classified as positive when it was observed at least three times across at least two follow-up years. Candidates with no future observations during the five-year window were classified as negative, while those that were observed without meeting the requirements for abundance and temporal breadth were classified as ambiguous.

The full *NoroScope* architecture contained 27 explicitly selected features distributed across five biological feature groups (Fig. 4, Table. S1). Eight ESM-2 features summarised candidate compatibility and variability across historical sequence contexts. Eight evolutionary features described both general site variability and the direct recurrence history of the candidate amino acid. Structural evidence was represented by two FoldX stability features and five Rosetta HBGA-binding features, while four binary annotations identified membership of major functional regions.

**Figure 4:**
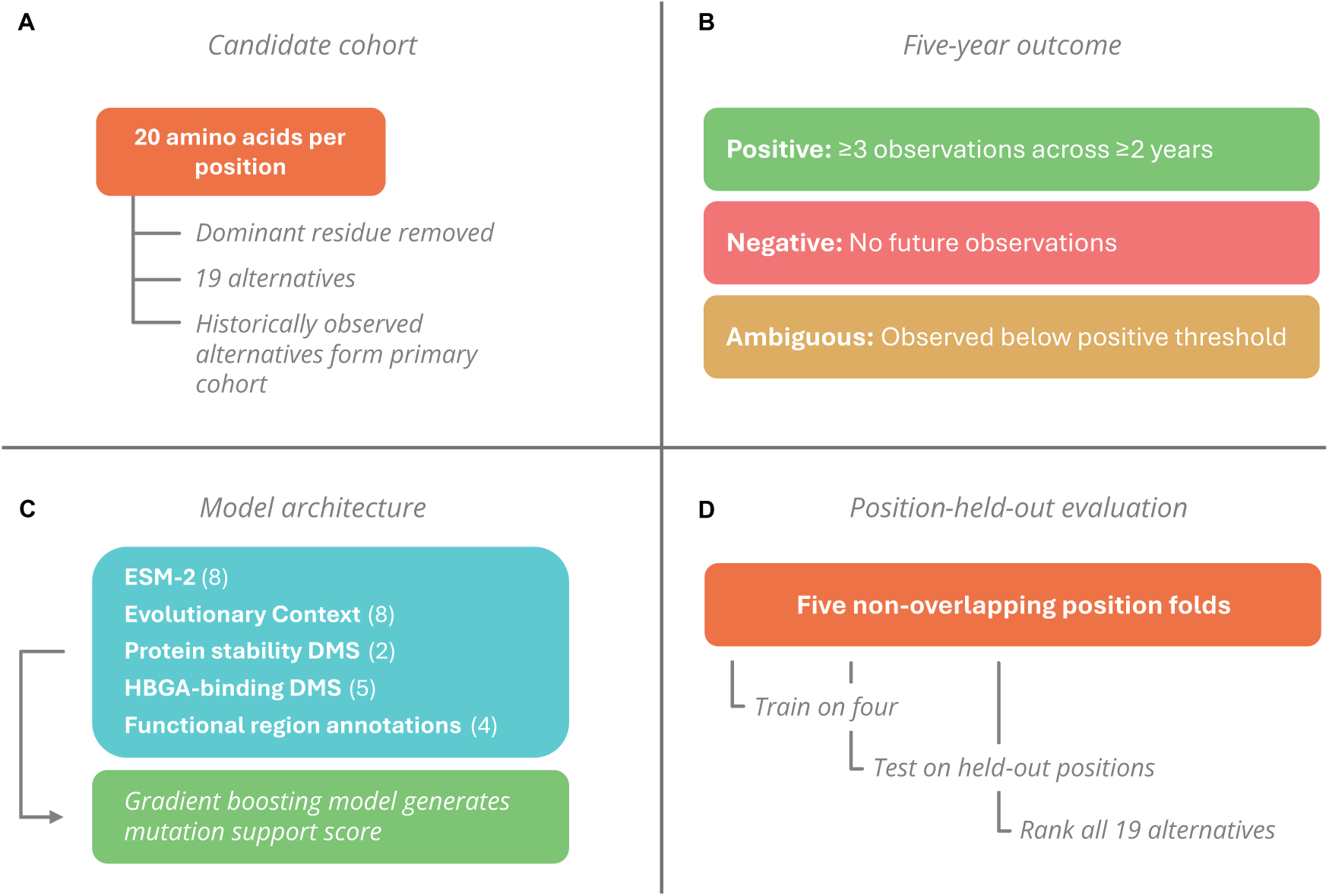
Construction and formulation of the NoroScope model. A: Amino-acid candidate cohorts used for model evaluation. At each position, the dominant residue was excluded and the remaining 19 amino-acid alternatives formed the complete within-position ranking pool. Previously observed non-dominant alternatives formed the primary modelling cohort. **B:** Model labels were defined from five-year post-cutoff outcomes. Candidates observed at least three times across at least two follow-up years were labelled positive, candidates with no future observations were labelled negative, and candidates observed below the positive threshold were labelled ambiguous. **C:** Twenty-seven input features were supplied to a histogram gradient-boosting model, which generated a relative mutation-support score. **D:** Position-held-out evaluation divided VP1 positions into five non-overlapping folds. Models were trained on four folds and evaluated on the remaining held-out positions until every eligible position had been tested exactly once.

The model was evaluated by dividing eligible positions into five non-overlapping folds using stratified group splitting. Predictions from the five test folds were then pooled into complete out-of-fold datasets, ensuring that every candidate received exactly one score from a model that had not been trained on any outcome from its VP1 position.

Evaluation was conducted at two complementary levels. Binary candidate classification compared high-confidence positives with candidates that were not observed during follow-up. Complete within-position evaluation then scored all 19 non-dominant amino acids at every eligible held-out position, including historically unseen and ambiguous candidates as competitors.

### *NoroScope* generalises mutation-support relationships to held-out VP1 positions

#### The complete NoroScope model prioritises future-supported alternatives at held-out positions

The complete NoroScope model strongly discriminated between high-confidence future-supported mutation candidates and alternatives that were not observed during follow-up (Fig. 5). Average precision was 0.805 at the 2011 cutoff and 0.834 at the 2016 cutoff. Receiver operating characteristic area under the curve (ROC-AUC) values were 0.856 and 0.904, respectively, showing that strong candidate discrimination was maintained across both historical settings.

**Figure 5:**
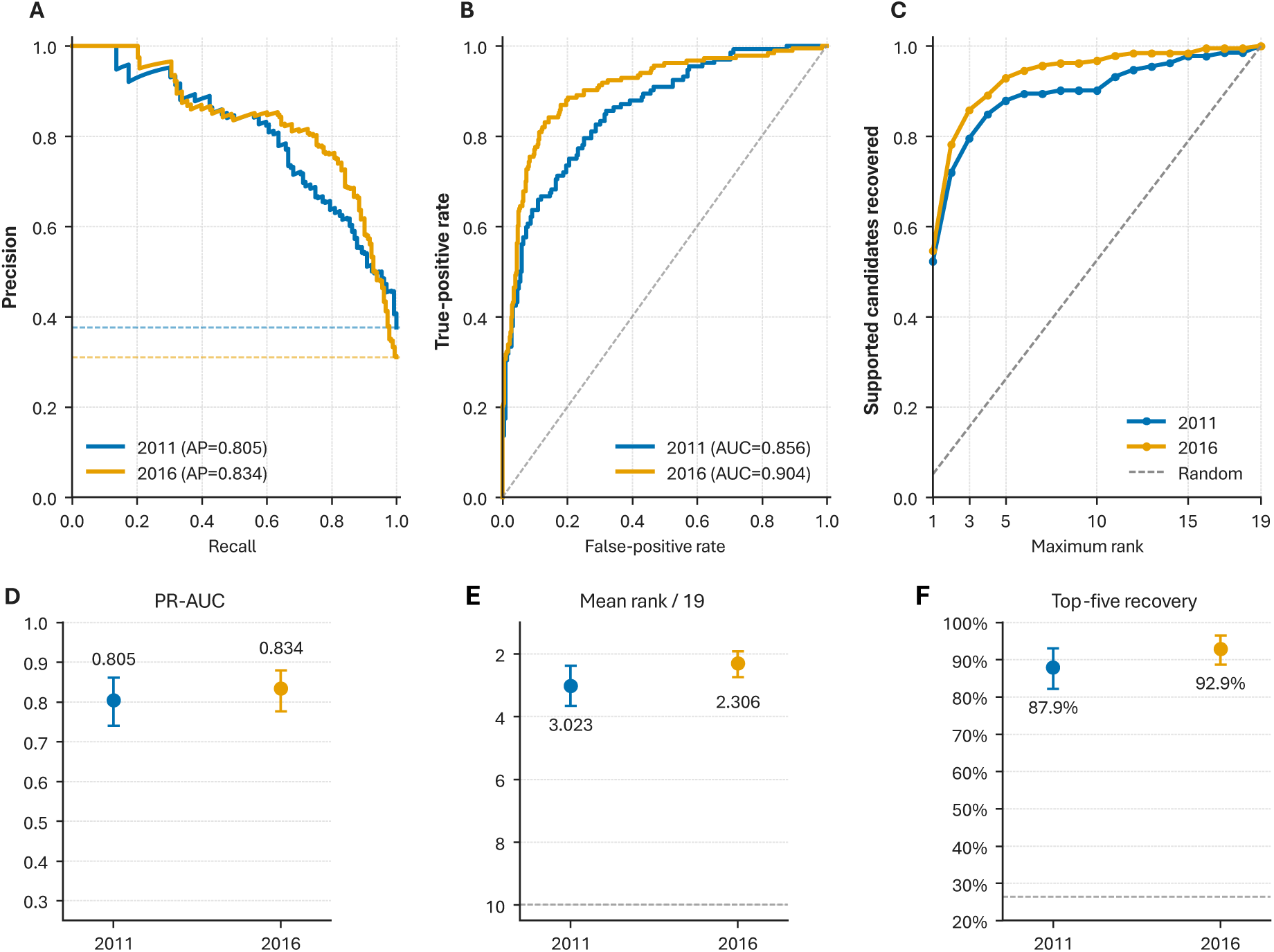
Performance of the complete NoroScope model at held-out VP1 positions. Performance of the complete 27-feature model was assessed using pooled five-fold position-held-out out-of-fold predictions for the 2011 and 2016 retrospective cutoffs. High-confidence future support was defined as at least three observations across at least two years during the five-year follow-up period. **A:** Precision–recall curves for classification of previously observed, non-dominant alternatives, with dashed line indicators of positive prevalence at each cutoff. **B:** Receiver operating characteristic curves, including an indicator of random discrimination. **C:** Cumulative recovery of future-supported candidates as the maximum accepted within-position rank was increased from 1 to 19. **D:** Precision-recall area under curve (PR-AUC) for the two cutoff-specific models. **E:** Mean within-position rank of future-supported candidates. Lower values indicate better prioritisation. **F:** Proportion of future-supported candidates ranked within the top five alternatives. Dashed lines indicate expected results from random scoring. Error bars indicate position-clustered 95% bootstrap confidence intervals.

The model’s strong global discrimination was also reflected in within-position ranking. Future-supported candidates received a mean rank of 3.02 among 19 alternatives in 2011 and 2.31 in 2016, substantially better than the expected mean rank of 10 under random ordering. NoroScope placed 87.9% and 92.9% of high-confidence future-supported candidates within the top five ranked alternatives in 2011 and 2016, respectively, compared with a random expectation of 26.3%.

#### Complementary feature blocks contribute distinct classification and ranking signals

Relative to a model trained only on site-level evolutionary context, the addition of candidate frequency, temporal breadth, age and recency improved mean within-position rank by 6.85 positions in 2011 and 7.96 positions in 2016 (Fig. 6). Candidate history was also strongly informative when added to molecular and site-level features, improving mean rank by a further 2.05 and 2.14 positions, respectively.

**Figure 6:**
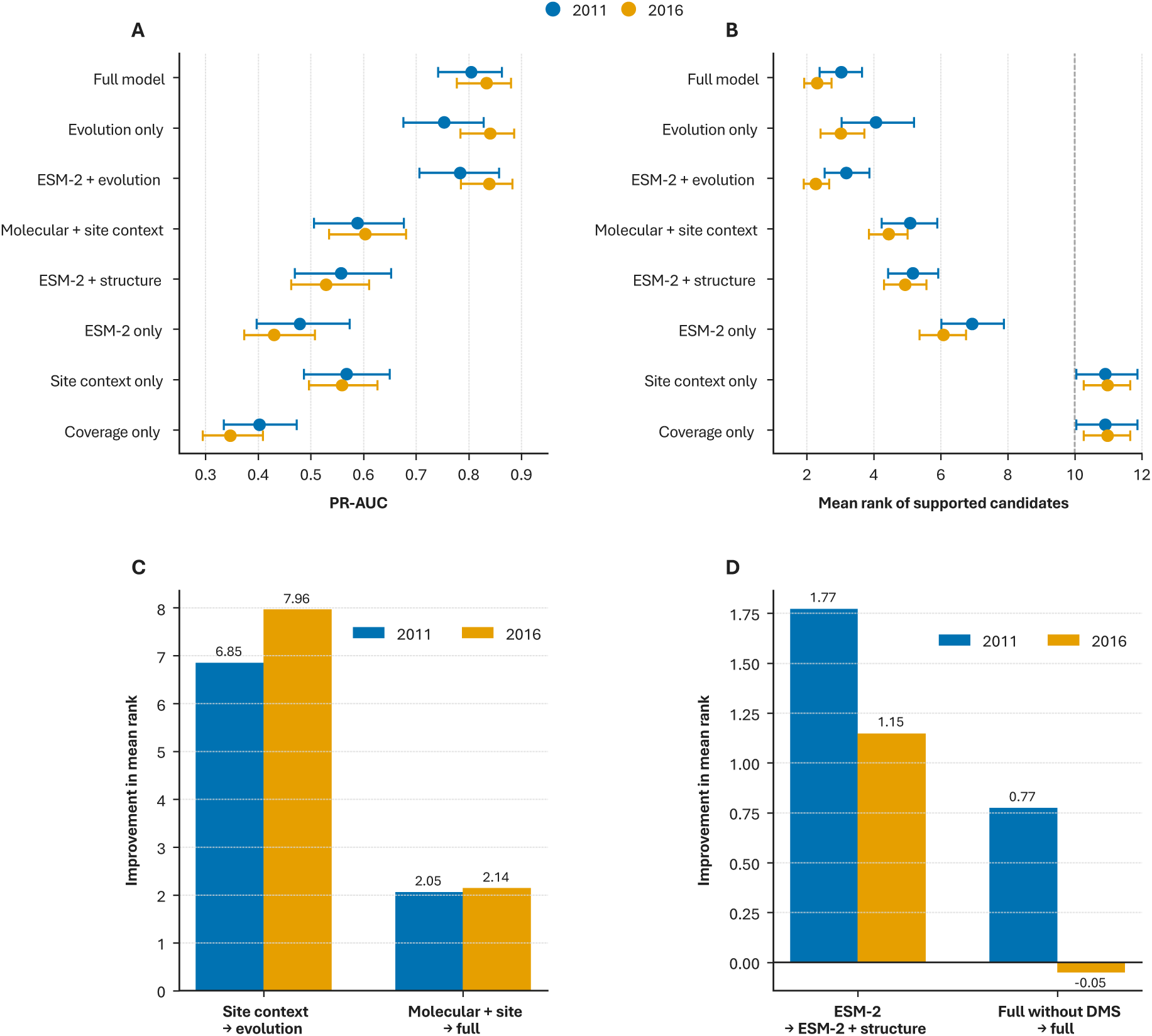
Feature ablation analysis of the NoroScope model. Performance was assessed using pooled five-fold position-held-out out-of-fold predictions for the 2011 and 2016 cutoffs. **A:** Precision-recall area under curve (PR-AUC) for predefined feature combinations in the primary observed-alternatives modelling cohort. The “molecular + site context” model combined ESM-2, DMS stability and binding features, functional annotations and site-level evolutionary features while excluding direct candidate-history variables. The coverage-only control used only indicators of ESM-2 and DMS feature availability. **B:** Mean within-position rank of future-supported candidates when all 19 non-dominant amino-acid alternatives were scored at each held-out position. Lower values indicate better prioritisation, and the dashed vertical line indicates the random expectation of rank 10. Panels A and B show pooled out-of-fold estimates, with error bars indicating position-clustered 95% bootstrap confidence intervals. **C:** Improvement in mean positive rank following addition of direct candidate-history features to either site-level evolutionary context alone or the combined molecular and site-context model. **D:** Improvement in mean positive rank following addition of structural DMS features to ESM-2 alone or to the otherwise complete model. Positive values indicate that the added feature block reduced, and therefore improved, mean positive rank.

The relative performance of the feature sets differed when evaluated within positions. Site-level evolutionary context retained moderate global discrimination but produced mean positive ranks close to random, indicating that it could identify positions enriched for future-supported mutations without reliably identifying the corresponding amino acid. In contrast, candidate-specific molecular features, including ESM-2 and structural information, retained within-position signal and substantially improved ranking among alternatives at the same site. The complete model performed best or close to best across both global discrimination and within-position recovery.

Structural information from the stability and binding DMS datasets provided a smaller but more targeted contribution. Adding structural and functional annotation features to a model trained only on ESM-2 improved mean positive rank by 1.77 positions in 2011 and 1.15 positions in 2016. When added to the otherwise complete model, DMS features improved mean rank by 0.77 positions in 2011 but produced no corresponding aggregate gain in 2016.

#### Model performance is not explained by positional shortcuts or feature coverage

To determine whether model performance arose from merely identifying variable positions rather than correctly prioritising supported amino acids, outcome labels were shuffled among candidates within each position (Fig. 7). Label shuffling substantially worsened mean rank and top-five recovery at both cutoffs, with performance falling close to what would be expected from random scoring across both global and within-position metrics.

**Figure 7:**
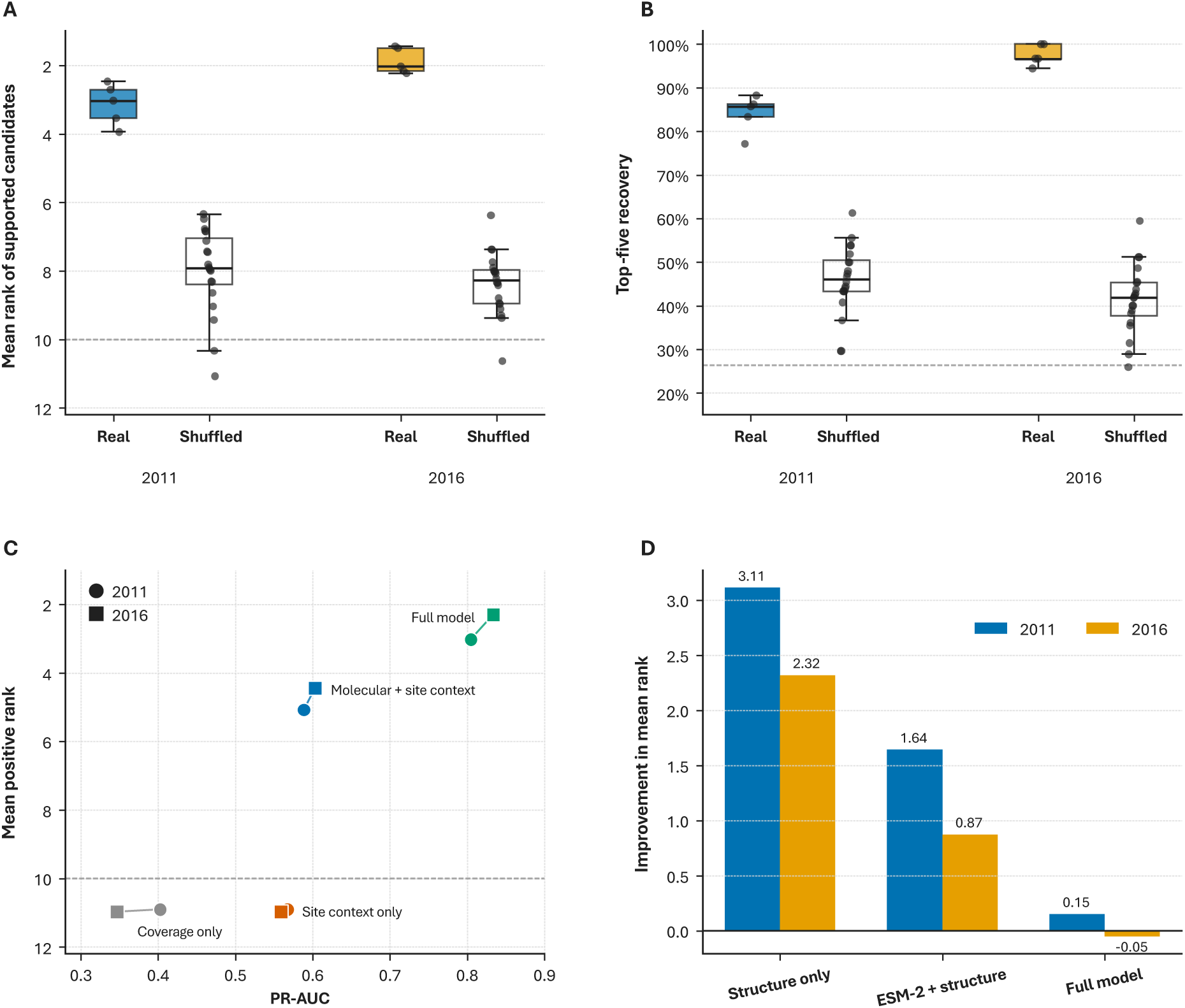
Control analyses for positional shortcuts and structural-feature coverage in NoroScope. A: Mean within-position rank of future-supported amino acid candidates using true outcome labels compared with a position-preserving label-shuffle control, in which labels were shuffled among candidates at the same VP1 position. Lower ranks indicate better prioritisation, and the dashed line marks the random expectation of rank 10. **B:** Corresponding recovery of future-supported candidates within the top five ranked alternatives. The dashed line indicates the random expectation of 26.3%. **C:** Relationship between global Precision-recall area under curve (PR-AUC) and mean within-position positive rank for the coverage-only, site-context-only, molecular-plus-site-context and complete models. Site-level and coverage-only models retained some global discrimination but ranked supported amino acids close to random, whereas candidate-specific features improved within-position recovery. **D:** Improvement in mean positive rank obtained from numerical FoldX and Rosetta values relative to a structural-missingness control that preserved the pattern of feature availability while replacing observed DMS values with constants. Positive values indicate improved ranking from the numerical structural measurements.

Models using only site context or feature-coverage indicators achieved some global discrimination but performed approximately at random in the within-position ranking task. Meanwhile, molecular features improved within-position ranking even without direct candidate history, and the complete model was able to combine strong candidate discrimination with accurate residue prioritisation.

A structural-missingness control was also used to test whether models incorporating DMS features benefited from the simple knowledge that structural coverage was restricted to the VP1 P domain. Replacing observed DMS values with constants while preserving their missingness pattern substantially reduced ranking performance for structure-only models, by 3.11 mean-rank positions in 2011 and 2.32 in 2016. Numerical DMS values also improved ESM-2-plus-structure models by 1.64 and 0.87 positions, respectively.

### Disagreement between evolutionary and molecular evidence reveals complementary mutation signal

To examine how these complementary evidence sources interacted with each other to influence individual predictions, we analysed a selection of positions within VP1 where the recurrence-only and molecular-only models agreed or disagreed when ranking each alternative amino acid. Here, the recurrence model used the complete evolutionary context feature set, while the molecular model excluded direct candidate history while including features from ESM-2, DMS, functional annotations and site-level evolutionary context.

Across VP1, most future-supported amino acid candidates were ranked within the top five by both the molecular and recurrence models: 73 of 132 candidates in 2011 and 113 of 183 in 2016 (Fig. 8). However, substantial disagreement remained. The recurrence model alone recovered 38 candidates in 2011 and 54 in 2016, corresponding to approximately 29% of supported candidates at each cutoff. Molecular evidence uniquely recovered a smaller but distinct group of 13 candidates in 2011 and 12 in 2016. Only eight and four supported candidates, respectively, were missed by both models.

**Figure 8:**
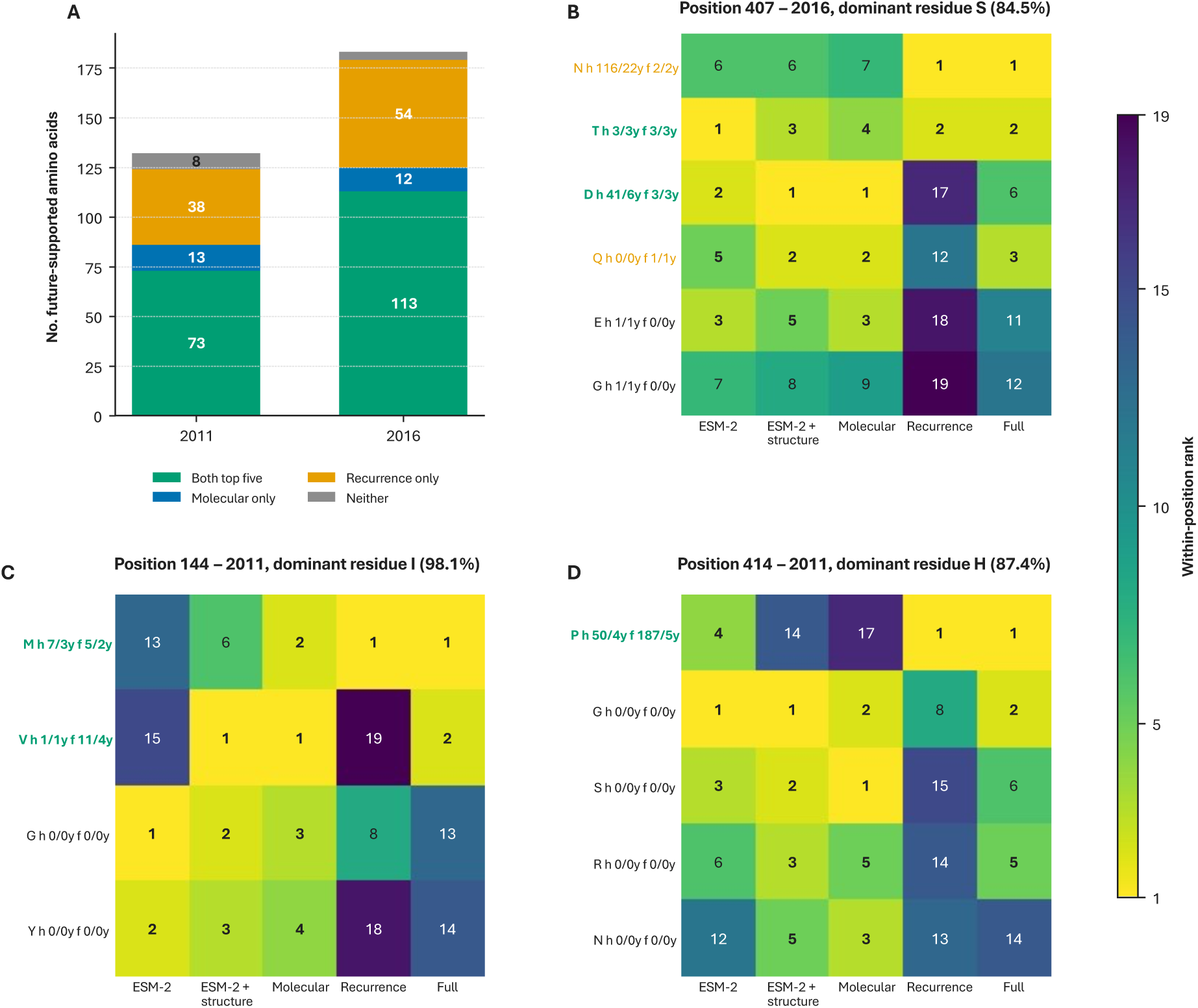
Disagreement between molecular and evolutionary evidence sources and representative position-level case studies. A: High-confidence future-supported candidates classified according to whether they were ranked within the top five alternatives by both the molecular and recurrence models, by either model alone, or by neither model. The molecular model combined ESM-2, structural DMS, functional-annotation and site-context features while excluding direct candidate-history variables. The recurrence model used the complete evolutionary-context dataset, including both site-level and candidate-history features. **B–D:** Within-position ranks assigned by the ESM-2-only, ESM-2-plus-structure, molecular, recurrence and complete models at representative VP1 positions: 407 at the 2016 cutoff (B), 144 at the 2011 cutoff (C) and 414 at the 2011 cutoff (D). Rank 1 indicates the highest-scoring candidate and rank 19 the lowest. The dominant amino acid at each position was excluded from the candidate pool, with its historical frequency shown above the heatmap. Candidate labels report historical observations and number of years represented before the cutoff (h n/y), followed by future observations and follow-up years represented (f n/y). The recurrence model can distinguish previously observed alternative amino acids using their candidate histories, but historically unseen alternatives lack these features and consequently receive equivalent scores within a position. Their individual recurrence ranks therefore reflect deterministic tie-breaking rather than candidate-specific evolutionary evidence.

At the 2011 cutoff, model scores for position 144 provided a clear example of molecular rescue. The methionine (M) alternative had already been observed across several historical sequences and years and was ranked first by the recurrence model. By contrast, valine (V) had only a single historical observation and was ranked last by recurrence despite subsequently receiving repeated support across four follow-up years. Molecular features ranked V first, and the complete model placed M and V first and second, respectively.

The reverse pattern occurred at position 414 in 2011. The proline (P) alternative had extensive historical support and was highly represented during follow-up, but was ranked poorly by the molecular model. Recurrence placed P first, and the complete model retained it as the top-ranked candidate.

Together, these results show that disagreement between the molecular and evolutionary evidence used by NoroScope can itself be informative. Such conflicts may identify positions where competing constraints are acting, where evolutionary pressure is elevated, or where important biological influences are not captured by the current feature set. These positions therefore represent particularly rich targets for subsequent experimental investigation and hypothesis generation.

### Experimental VLP assembly supports contrasting molecular predictions at position 297

Position 297 was initially selected for experimental investigation following zero-shot ESM-2 analysis of the NoroScope sequence dataset, before construction of the final integrated NoroScope model. Asparagine (N) received an unusual highly positive ESM-2 LLR despite having only two observations in the sequence dataset, both in 2019. By contrast, tryptophan (W) received the lowest ESM-2 LLR at the position, despite also having two observations in the dataset.

The strong contrast between the two substitutions was maintained after the construction of the final NoroScope framework (Fig. 9). Across the 2011 and 2016 historical cutoffs, the mean raw ESM-2 probability assigned to 297N was 71.2% and 68.5%, respectively, compared with approximately 0.001% for 297W. The complete model consistently placed N above W and among the more highly ranked alternatives at position 297.

**Figure 9:**
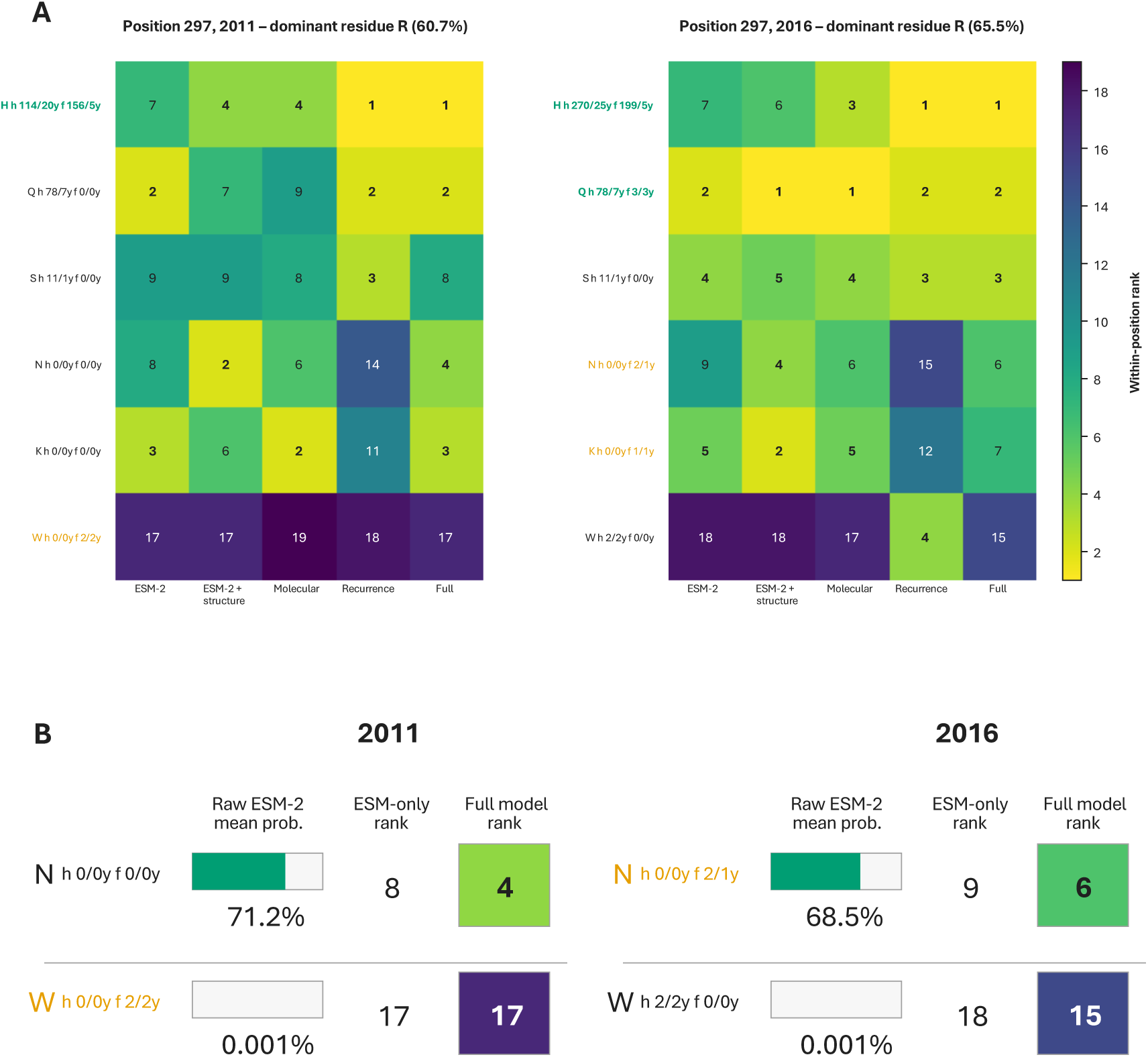
Comparison of experimental-validation candidates 297N and 297W across ESM-2 and NoroScope predictions. A: Within-position ranks for selected amino-acid alternatives at VP1 position 297 under the 2011 and 2016 historical cutoffs. Candidates are compared across the ESM-2-only, ESM-2-plus-structure, molecular-plus-site-context, recurrence-only and complete NoroScope models. The dominant residue, R, was excluded from ranking. Labels indicate the number of historical observations and years represented before each cutoff (h n/y) and the corresponding values within the subsequent five-year evaluation window (f n/y). **B:** Direct comparison of raw zero-shot ESM-2 mean candidate probabilities with ESM-2-only and complete model within-position ranks for 297N and 297W. Historically unseen alternative amino acids lack candidate-specific recurrence information and therefore receive equivalent recurrence model scores within a position. Differences in their displayed recurrence ranks reflect deterministic tie-breaking rather than meaningful discrimination between unseen candidates.

To test whether the contrasting ESM-2 and NoroScope predictions for 297N and 297W corresponded to changes in the ability of VP1 to self-assemble into VLPs, each substitution was engineered independently into an established baculovirus expression plasmid containing the GII.4 open reading frame 2 (ORF2, coding for VP1) sequence from a virus collected in 2004 (28). Recombinant VP1 was expressed using the baculovirus system, and the formation of VLPs was visualised using transmission electron microscopy (TEM).

TEM revealed that 297N formed abundant structures with morphology consistent with HuNoV VLPs, indicating that the substitution was compatible with VP1 self-assembly (Fig. 10). No comparable particle formation was observed for 297W under the same conditions.

**Figure 10:**
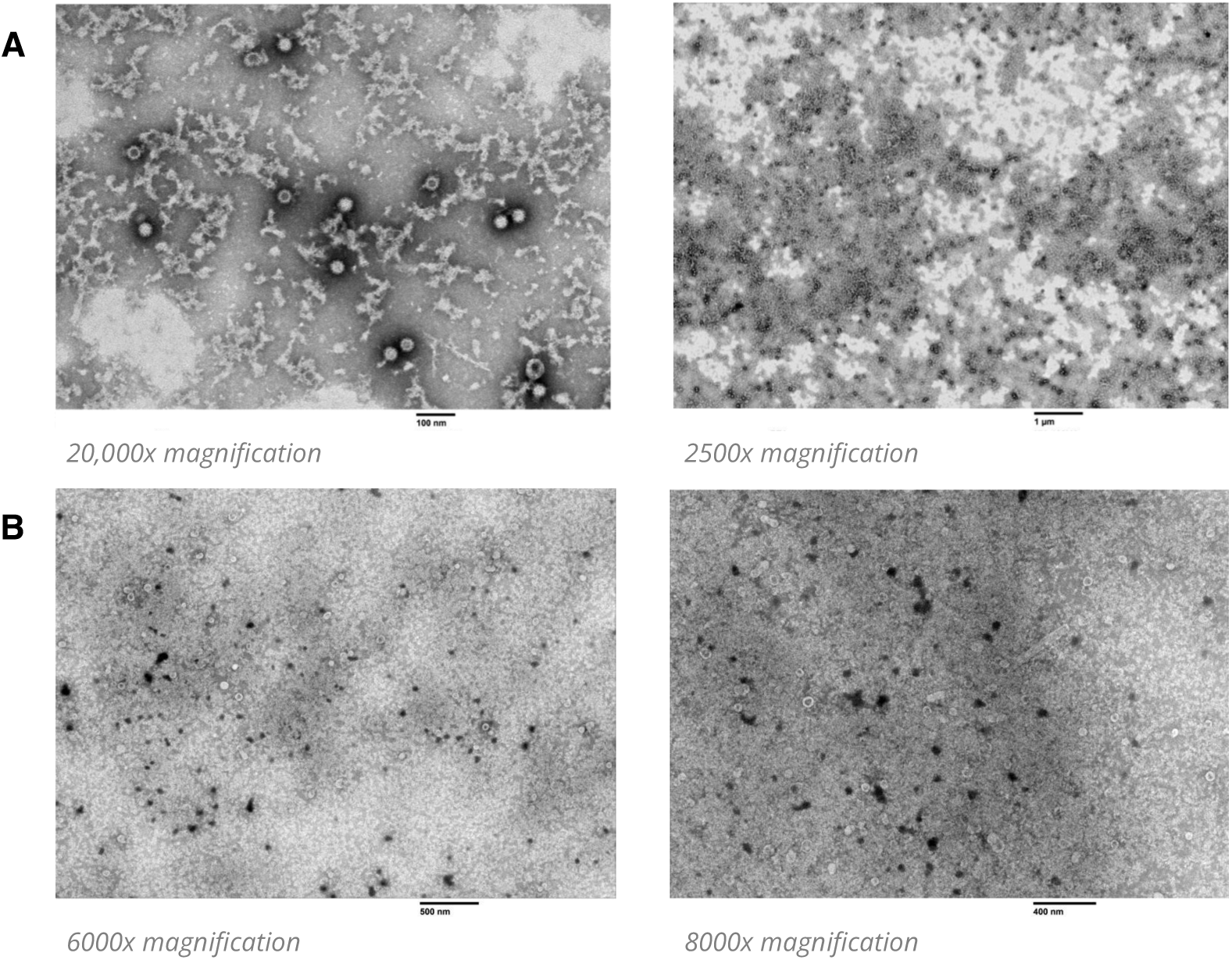
Transmission electron micrographs of recombinant VP1 carrying NoroScope validation substitutions at position 297. A: Representative micrographs following expression of VP1 containing the 297N substitution, showing abundant structures with morphology consistent with virus-like particles. **B:** Representative micrographs following expression of VP1 containing 297W, for which no comparable particle formation was observed under the tested expression conditions. Image magnifications and scale bars are shown for each panel.

Together, these contrasting assembly phenotypes were consistent with the strong difference in molecular support assigned to 297N and 297W by ESM-2 and retained within the integrated NoroScope model. Although restricted to a single position and sequence background, the experiment nevertheless provides independent evidence that the contrasting molecular predictions at position 297 reflect a biologically meaningful difference in VP1 assembly.

## Methods

### Construction of the GII.4 VP1 sequence dataset

The principal longitudinal sequence dataset used by NoroScope was derived from the NOROPATROL collection of HuNoV genomes obtained from clinical samples collected in England and Wales between 1994-2023 (16). Candidate ORF2 sequences were identified using ORFipy v0.0.4 (29). Capsid genotype information was used to identify GII.4 sequences, with genotype assignments independently checked using the RIVM norovirus typing tool (30).

Additional HuNoV sequences were obtained from NCBI Virus/GenBank to increase the geographical and temporal representation contained within the model dataset (31). ORF2 extraction was performed using the same ORFipy parameters used for the NOROPATROL sequences.

The GenBank-derived and NOROPATROL-derived GII.4 VP1 sequences were combined using exact protein-sequence matching. The resulting protein sequences were aligned using MAFFT with automatic algorithm selection (--auto) and otherwise default parameters (32).

Post-alignment quality control was performed relative to the GII.4/2002 v2 VP1 reference sequence described by Allen et al. (28). Alignment columns corresponding to reference-specific gaps were removed to establish a common 540-position VP1 coordinate system. Sequences containing non-canonical insertions or deletions relative to this coordinate system were excluded, while the lineage-associated single-residue gap corresponding to NoroScope position 394 was explicitly permitted.

The final NoroScope modelling dataset comprised 1,657 unique, dated GII.4 VP1 sequences, including 1,009 sequences originating from GenBank and 648 from NOROPATROL, spanning sample collection years from 1974 to 2024.

Functional regions within GII.4 VP1 were defined before model analysis using published structural, antigenic and population-genomic studies. Three broad functional categories were represented: antigenic epitopes, the HBGA binding pocket and the conserved conformational “breathing core”.

### ESM-2 zero-shot mutation scoring

ESM-2 was used to estimate the sequence-context compatibility of alternative amino acids at each VP1 position using zero-shot masked language modelling (6,33). Four pretrained ESM-2 architectures were initially compared: the 8-million-, 650-million-, 3-billion- and 15-billion-parameter models. The 15-billion-parameter ESM-2 model produced the strongest overall performance across these comparisons and was therefore selected for the full NoroScope dataset.

Zero-shot mutation scores for the complete sequence dataset were generated using the pretrained 15-billion-parameter ESM-2 checkpoint facebook/esm2_t48_15B_UR50D (6). The scoring implementation was adapted from a Hugging Face ESM-2 mutation-scoring implementation, with modifications for batched dataset-scale inference and the NoroScope analysis workflow (34).

For each VP1 sequence, every position containing a standard amino acid was masked individually while the remainder of the sequence was retained as context. ESM-2 then generated a probability distribution across the 20 canonical amino acids at the masked position. For candidate amino acid a at position *i*, a log-likelihood ratio (LLR) was calculated relative to the amino acid observed in the input sequence:

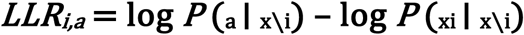

Positive values indicate that the candidate received greater model support than the observed residue within that sequence context, whereas negative values indicate lower relative support.

### Structural preparation and *in silico* deep mutational scanning

Experimental X-ray crystallographic structures of HuNoV GII.4 VP1 P-domain dimers were obtained from the Protein Data Bank (35). For the protein-stability scan, three ligand-free P-domain structures were used: 4OOV, 4OOX and 5IYN. For the HBGA binding scan, structures 4WZT, 4X07 and 5IYP were selected, each containing a GII.4 P-domain dimer in complex with a type A HBGA glycan.

Each experimental template was then remodelled to represent four GII.4 VP1 sequence backgrounds used in the experimental component of the study: GII.4/1999 v0 and GII.4/2002 v2, described previously by Allen et al., and GII.4/2009 and GII.4/2012, corresponding to GenBank accessions MZ376650.1 and MZ376651.1, respectively (28,36).

Mutation-associated effects on VP1 protein stability were estimated using FoldX and automated via the MutateX workflow (26,37). Saturation mutagenesis was performed across all VP1 positions with structural representations. For each substitution, FoldX estimated the change in folding free energy relative to the corresponding wild-type structure:

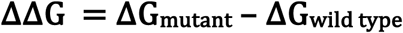

Positive ΔΔG values therefore indicated predicted destabilisation, whereas negative values indicated predicted stabilisation. One structure–sequence input, corresponding to the 5IYN-derived v0 model, behaved as a major technical outlier and was excluded from the final stability dataset.

Mutation-associated effects on the interaction between VP1 and the bound HBGA were estimated using a paired wild-type–mutant workflow implemented in Rosetta (27). Binding-associated energy was calculated using Rosetta InterfaceAnalyzer. Mutation-associated binding effects were defined from paired calculations as:

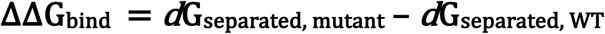

Positive ΔΔGbind values indicated predicted weakening of the VP1–HBGA complex relative to the paired wild-type calculation, whereas negative values indicated more favourable predicted interface energetics.

Repeated paired calculations were performed to reduce stochastic variation and assess convergence of each mutation-associated estimate. An initial minimum of 10 paired replicates was generated for each mutation–structure–sequence combination. Combinations with a standard error of the mean greater than 0.25 Rosetta energy units were expanded in batches of 10 paired replicates, to a maximum of 30. (fig. S3)

FoldX and Rosetta calculations were treated separately during the data preparation stage rather than being folded into a single value.

### Construction of the temporally-controlled NoroScope database

The NoroScope sequence dataset, ESM-2-derived LLRs, structural DMS results and functional-annotation data were integrated within a relational database implemented using DuckDB (38). All datasets were mapped to the common 540-position NoroScope VP1 coordinate system and linked at the level of sequence background, VP1 position and candidate amino acid.

FoldX and Rosetta measurements were available for the four representative VLP sequence backgrounds rather than independently for every surveillance sequence. Each surveillance sequence was therefore assigned the most appropriate structural background before DMS features were linked. Positions outside the structurally represented P domain retained missing DMS values rather than being assigned a neutral structural effect.

Evolutionary-context features were calculated independently at each historical cutoff using only dated VP1 sequences sampled on or before that year. At the positional level, amino-acid variability was represented by Shannon entropy, together with the number of distinct amino acids observed and the frequency of the dominant residue. Candidate-specific evolutionary history was represented by candidate frequency, the number of distinct sampling years in which the candidate had been observed, candidate age and candidate recency.

The years 2011 and 2016 were selected as retrospective prediction cutoffs. Historical feature construction and future outcome construction were implemented as separate database stages. Sequence observations after each cutoff were excluded from all molecular and evolutionary model features and were accessed only when constructing future-support outcomes.

### Definition of the mutation prediction task and future support outcomes

For each historical cutoff and VP1 position, the amino acid with the highest observed frequency among sequences available to the model was designated the dominant residue. The dominant residue was excluded from the mutation-prediction task, leaving the remaining 19 canonical amino acids as the complete alternative-candidate pool at each position. The primary modelling cohort comprised historically observed, non-dominant alternatives.

The five-year window was used for the primary NoroScope analysis. A candidate amino acid was classified as positive when it was observed in at least three post-cutoff sequences distributed across at least two distinct follow-up years. Candidates receiving no observations during the five-year period were classified as negative. Candidates that were observed during follow-up but did not satisfy both the abundance and temporal-breadth criteria were classified as ambiguous. Ambiguous candidates were excluded from binary model fitting and from calculation of global classification metrics. During complete within-position ranking, however, ambiguous candidates remained among the 19 competing alternative amino acids.

Separate models were fitted for the 2011 and 2016 historical cutoffs. The complete NoroScope model contained 27 predefined features representing complementary sources of molecular and evolutionary information. These comprised eight ESM-2 sequence-context features, four site-level evolutionary features, four candidate-history features, two FoldX stability features, five Rosetta HBGA-binding features and four functional-region annotations.

Models were fitted using the HistGradientBoostingClassifier implemented in scikit-learn (39). No negative-class downsampling was performed; instead, balanced sample weights were calculated from the positive and negative observations within each training partition.

The primary model evaluation used five-fold position-grouped cross-validation. All candidates associated with a given position were therefore assigned to the same fold. Consequently, every candidate in the primary binary cohort was scored once by a model that had not been trained using outcomes from its position.

Global discrimination was assessed using average precision and ROC-AUC. Candidate-specific performance was assessed through within-position ranking. The principal ranking metrics reported were the mean rank of high-confidence future-supported candidates and the proportion of these candidates appearing within the top five alternatives.

The contribution of individual evidence sources was evaluated using predefined feature subsets fitted using the same candidate cohorts and validation framework. Three principal controls were used to test whether model performance could arise from non-candidate-specific shortcuts. These comprised a coverage-only model, a position-preserving label-shuffle control and a structural-missingness control.

### Experimental validation of VP1 assembly

VP1 position 297 was selected for experimental investigation during zero-shot ESM-2 analysis, before construction of the final integrated NoroScope model. The R297N and R297W substitutions were introduced independently into a pRN16-based expression plasmid containing ORF2 from the GII.4/2004 v2 VP1 background. The underlying GII.4 VLP expression constructs were derived from the recombinant baculovirus system described previously by Allen et al. (28).

Mutant plasmids were generated using the Q5 Site-Directed Mutagenesis Kit (New England Biolabs) and verified by whole-plasmid sequencing. Recombinant baculovirus was generated by co-transfection of Sf9 cells with BAC10:KO1629 bacmid DNA and the corresponding mutant VP1 expression plasmid. Recombinant viruses were purified by plaque assay.

For preparative VP1 expression, Sf9 suspension cultures were infected at a multiplicity of infection of 2–3 and incubated for 48–72 h at 28°C. VP1-containing material was purified by ultracentrifugation through a discontinuous 15%/60% (w/v) sucrose cushion.

Purified VP1 preparations were analysed by transmission electron microscopy (TEM) to assess formation of structures morphologically consistent with HuNoV VLPs. TEM sample preparation and imaging were performed by Dr Ana Godi at the UK Health Security Agency (UKHSA). R297N and R297W preparations were processed using the same procedure for qualitative comparison of particle assembly.

## Discussion

In this study, we developed NoroScope as a transferable, ML-based framework for investigating the mutational landscape of viral proteins, with a particular focus on targets for which comparatively limited surveillance and experimental data are available. NoroScope sought to make greater use of the available viral sequence data for HuNoV GII.4 by supplementing it with complementary biological information derived from ESM-2 and structural modelling alongside evolutionary history. By retaining each source as a distinct feature block, NoroScope was designed not only to prioritise plausible amino-acid alternatives at different positions, but also to investigate what each source of biological information contributes and where their integration provides additional value.

Overall, results support this central premise. Across both retrospective cutoff years of 2011 and 2016, the complete NoroScope model strongly distinguished future-supported from unsupported alternative amino acids and consistently prioritised supported residues near the top of its within-position rankings. The integrated model performed best or close to best across the principal evaluation tasks, while ablation and control analyses showed that this performance could not be explained simply by identifying mutation-prone positions or patterns of structural-data availability. Although the different feature sources contributed unequally to overall predictive performance, each captured distinct aspects of mutation plausibility.

To begin, the ESM-2-derived feature set was constructed to directly address one of this study’s central questions: Can foundation pLLMs be used to provide additional useful biological information for viruses with otherwise limited data? When considered independently, ESM-2-derived features retained substantial candidate-specific predictive signal, ranking future-supported amino-acid substitutions considerably better than random across both historical cutoffs.

Beyond its predictive performance, several results support the biological relevance of the information transferred by ESM-2 to GII.4 VP1. Most notably, the strongly contrasting predictions for 297N and 297W were consistent with their differing VLP-assembly phenotypes.

At the same time, improvements in model performance observed following incorporation of target-specific structural and functional information show that these pLLM representations do not capture every constraint relevant to VP1 mutation. Instead, ESM-2 provides a broad and transferable molecular prior, while the integrated NoroScope model gains additional value by combining this prior with explicit structural and evolutionary context specific to GII.4 VP1.

In contrast to the broad sequence representation provided by ESM-2, the structural feature sets relating to results from DMS studies provide explicit estimates of particular molecular constraints acting on GII.4 VP1. FoldX-derived scores describe predicted mutation-associated effects on protein stability, while Rosetta-derived scores describe effects on the energetics of the modelled VP1–HBGA complex. These structural features retained genuine candidate-specific signal, with combined FoldX and Rosetta models ranking future-supported substitutions above chance. Furthermore, the near absence of correlation between FoldX and Rosetta scores suggests that the two approaches capture distinct aspects of the structural constraints acting upon VP1, rather than alternative measurements of the same underlying property. Adding structural and functional-annotation features to a model trained otherwise only on ESM-2 improved mean within-position rank from 6.93 to 5.16 in 2011 and from 6.08 to 4.93 in 2016.

Lastly, the evolutionary context dataset provides an explicit representation of how GII.4 VP1 has varied through time. These features, which captured knowledge of which amino-acid alternatives had previously occurred, how frequently they had been observed and how broadly those observations were distributed through time, represented the strongest individual source of predictive information within the model. This is perhaps unsurprising, given that the ECD features provide information to the model which is directly relevant to its prediction task, namely the future support of previously observed alternative amino acids. In contrast, information captured by the ESM-2 and structural feature sets provides more indirect measurements of molecular plausibility. However, the strength of the ECD also defines an important limitation of evolutionary history as a predictive signal. For amino acids that have never previously been observed at a given position, direct candidate history features contain no information with which to distinguish one unseen alternative from another. Molecular features, then, provide an important complementary source of candidate-specific evidence, allowing the model to leverage each feature set according to the information available and generalise learned relationships across the protein even where particular forms of evidence are sparse or absent.

The differing contributions of the NoroScope feature sets also provide an opportunity to examine how agreement and disagreement between them can be interpreted biologically. These evidence sources are related, but they are not expected to agree in every case. Natural recurrence depends on molecular compatibility alongside additional factors including host immunity, transmission, genetic background, stochastic processes and epidemiological opportunity. A substitution may therefore appear molecularly plausible without becoming recurrent in nature, just as a naturally successful substitution may depend on biological pressures or sequence-context effects that are not fully represented by the current molecular features.

Across VP1, the molecular and recurrence feature sets agreed more often than they disagreed, with both models placing the majority of future-supported amino acids within their top five alternatives at each cutoff. Nevertheless, molecular predictions recovered a distinct subset of candidates missed by recurrence, further demonstrating their contribution of complementary candidate-specific information beyond that provided by evolutionary history alone. Position 144 provides a particularly clear example of molecular features rescuing sparse recurrence data, while position 414 shows the reverse situation: the strongly recurrent proline alternative was ranked poorly by the molecular model but restored to first place by evolutionary history, suggesting that important constraints or selective pressures at this position are not fully represented by the current molecular feature set. Such scenarios provide ideal targets for further investigation and experimental validation, supporting a “lab in the loop” workflow that sees experimental results used to iterate upon and subsequently improve model predictions.

NoroScope is best interpreted as a framework for exploring and prioritising the mutation space available at individual positions within a viral protein. By integrating molecular and evolutionary evidence, it generates relative rankings of alternative amino acids and provides a structured means of identifying candidates supported by different forms of biological context. These rankings should be interpreted as relative mutation-support scores rather than calibrated probabilities of viability, fitness or future emergence. This distinction is important considering that the model prediction task is defined by observed recurrence in genomic surveillance data – while such data provide valuable evidence of observed amino acid states in natural populations, it is not comprehensive, and the presence or indeed absence of a given substitution from the surveillance record does not necessarily imply biological compatibility or lack thereof. We sought to reduce the influence of these uncertainties by defining future support conservatively, requiring repeated observations across multiple follow-up years for positive classification, while treating intermediate cases as ambiguous rather than forcing them into either class. While this strategy yielded insightful results, it was unable to overcome another limitation relating to the sequence dataset, that being a scarcity of historically unseen substitutions that subsequently receive future support within the NoroScope sequence dataset. This led to our selection of future support of previously observed amino acids as the primary prediction task, as too few positive examples of these unseen substitutions existed for reliable model training.

The structural feature sets introduce a separate set of limitations arising from the availability of suitable experimental VP1 structures and the approximations inherent to *in silico* DMS. FoldX and Rosetta calculations were restricted to the structurally represented P domain and generated across a limited number of representative structural templates and sequence backgrounds. We sought to assess the robustness of these estimates by comparing predictions across multiple experimental structures and VP1 backgrounds, and found that the *in silico* DMS datasets were broadly robust to variation in both structural template and sequence background, contributing useful biological information despite lacking unique representations for every individual sequence and being restricted to the P domain.

The natural next step for NoroScope is to expand the model sequence dataset beyond GII.4, incorporating additional HuNoV genotypes and broadening the experimental evidence available for model evaluation. We anticipate that this will provide a richer training and validation framework, allowing NoroScope to move from retrospective benchmarking towards genuinely prospective mutation prediction and targeted experimental testing. Beyond HuNoV, NoroScope provides a generalisable framework for integrating pretrained protein representations with target-specific molecular and evolutionary context. This approach can be transferred to other viral proteins and pathogens where sequence data are available but comprehensive experimental training datasets remain limited.

Taken together, this study defines NoroScope as a generalisable modelling framework for investigating the plausibility of amino acid substitutions in proteins. We show that pretrained protein representations can contribute biologically useful information to comparatively data-limited viral systems, and that this information can be strengthened through integration with target-specific structural and evolutionary context. By integrating these complementary signals, NoroScope provides a practical framework for mutation prioritisation, hypothesis generation and functional investigation in proteins.

