## Supplementary Information for "NoroScope: Exploring the Mutational Landscape of the Human Norovirus Capsid Protein with Context-Aware Machine Learning"

| Feature block | Feature | Description | Information represented |
|---|---|---|---|
| **ESM-2** | Median candidate LLR | Median ESM-2 log-likelihood ratio for the candidate amino acid across unique pre-cutoff VP1 sequence contexts. | Describes the typical ESM-2-derived LLR for the candidate amino acid at a position relative to the observed wild type. |
| **ESM-2** | Candidate LLR SD | Standard deviation of candidate amino acid LLR across unique pre-cutoff sequence contexts. | Describes the degree to which support for the chosen amino acid candidate is dependent on sequence context. |
| **ESM-2** | Candidate LLR 90th percentile | LLR value exceeded within only the highest-scoring 10% of sequence contexts. | Describes the level of ESM-2 support achieved by the candidate amino acid in only its most favourable sequence backgrounds. |
| **ESM-2** | Mean candidate rank | Mean rank of the candidate among the 19 non-wild-type alternatives within each sequence context. Rank 1 indicates the highest LLR. | Describes ESM-2 preference for the amino acid candidate compared to other residues at the same position. |
| **ESM-2** | Mean candidate percentile | Mean percentile rank among the 19 non-wild-type alternative amino acids at a position. Larger values indicate stronger relative ESM-2 support. | Provides a scale-independent measure of ESM-2 preference for the amino acid candidate compared to other residues at the same position. |
| **ESM-2** | Mean distance from best alternative | Mean difference between the highest alternative amino acid LLR and the candidate amino acid LLR within each sequence context. | Measures how far below the highest-ranked alternative amino acid at that position ESM-2 ranks the candidate. A value of 0 indicates that the candidate is the highest-ranked. |
| **ESM-2** | Median candidate probability | Median candidate amino acid probability at a position across all pre-cutoff sequence contexts, obtained by normalising ESM-2 LLRs across the 20 canonical amino acids. | Measures typical ESM-2 support for a given amino acid candidate at a position across all pre-cutoff sequence contexts. |
| **ESM-2** | Mean ESM-2 site entropy | Mean Shannon entropy of the ESM-2 20-amino-acid probability distribution across pre-cutoff sequence contexts. | Overall variability of a position, accounting for both number and frequency of observed amino acids. |
| **Evolutionary context – site** | Site entropy | Shannon entropy of observed amino-acid frequencies at the position using sequences sampled on or before the cutoff. | Describes historical amino acid diversity at a position. |
| **Evolutionary context – site** | No. residues observed | Number of unique amino acids observed at the position up to the cutoff date. | Describes the historical variety of amino acids appearing at a position. |
| **Evolutionary context – site** | Dominant residue frequency | Frequency of the most common amino acid at the position by the cutoff. | Describes how much a position has historically been dominated by a single amino acid. |
| **Evolutionary context – site** | No. distinct WT residues | Number of unique historical wild-type residues from which the candidate amino acid was represented as an alternative across available pre-cutoff sequence contexts. | Describes the number of unique wild-type amino acids that a candidate amino acid is compared against. |
| **Evolutionary context – candidate history** | Candidate frequency | Number of pre-cutoff sequence contexts containing the candidate amino acid divided by the total number of pre-cutoff sequence contexts which contain a residue. | Describes the broad historical commonality of the candidate amino acid at a position. |
| **Evolutionary context – candidate history** | Candidate years observed | Number of distinct pre-cutoff sampling years in which the candidate was observed. | Describes the broad temporal prevalence of the candidate amino acid at a position. |
| **Evolutionary context – candidate history** | Candidate age | Cutoff year minus the year that a candidate amino acid was first observed. Undefined if the candidate had not previously been observed. | Defines the length of the candidate amino acid's recorded evolutionary history within the NoroScope dataset within a position. |
| **Evolutionary context – candidate history** | Candidate recency | Cutoff year minus the year that a candidate amino acid was most recently observed. Undefined if the candidate had not previously been observed. | Informs the model on how recently the candidate amino acid has been observed relative to the cutoff year within a position. |
| **FoldX stability** | Median FoldX ΔΔG | Median capped FoldX ΔΔG across represented pre-cutoff structural background. Values were capped at −3 and +10 kcal $mol^{-1}$ before integration into the model dataset. | Describes the typical predicted effect of the candidate amino acid on P domain stability at a position. |
| **FoldX stability** | FoldX ΔΔG SD | Standard deviation of capped FoldX ΔΔG across represented pre-cutoff structural backgrounds. | Describes the extent to which the candidate amino acid's effect on P domain stability is influenced by sequence and structural context at a position. |
| **Rosetta HBGA binding** | Median binding ΔΔG | Median Rosetta consensus ΔΔGbind across represented pre-cutoff structural backgrounds. Template-consensus values were globally winsorised at the 1st and 99th percentiles before integration into the model dataset. | Describes the typical effect of the candidate amino acid on the binding energetics of the bound VP1 P domain-HBGA complex at a position. |
| **Rosetta HBGA binding** | Median position-centred binding residual | Median position-centred residual across represented pre-cutoff structural backgrounds. Residuals were calculated fro consensus scores which had not been winsorised. | Describes how a candidate amino acid affects HBGA binding at a position relative to the broad mutation sensitivity of the position itself. |
| **Rosetta HBGA binding** | Binding-residual SD | Standard deviation of position-centred residuals across represented pre-cutoff structural backgrounds. | Describes to what degree candidate-specific binding effects are dependent on sequence and structural context. |
| **Rosetta HBGA binding** | Mean binding QC weight | A mean 0-1 score of confidence in and quality of the binding estimate. Incorporates template availability and convergence status of substitutions. | Describes the overall reliability and completeness of Rosetta binding estimates for the candidate amino acid at a position. |
| **Rosetta HBGA binding** | Mean template agreement | Mean fraction of experimental templates giving the same ΔΔGbind direction within each represented pre-cutoff structural background. | Describes how consistently the different experimental templates for binding estimates predict the direction of the binding effect for the candidate amino acid at a position. |
| **Functional annotation** | Breathing-core membership | Binary indicator for positions 234, 310, 316, 484 or 493. | Confers membership of the "breathing core" functional region. |
| **Functional annotation** | HBGA-pocket membership | Binary indicator for membership of annotated HBGA loops 1–3 or the adjacent β-sheet. | Confers membership of the HBGA binding pocket functional region. |
| **Functional annotation** | Epitope membership | Binary indicator for membership of any annotated epitope A–I. | Confers membership of an antigenic region. |
| **Functional annotation** | Functional region of interest | Binary indicator for membership of any annotated epitope, the HBGA-binding pocket or the breathing core. | Confers membership of any labelled functional region. |

**Table S1: The complete NoroScope feature set.** Includes all mathematical features provided to the full NoroScope model during training. Features are divided into blocks describing the data source from which they originate, and each feature is accompanied by a description and explanation of the information it conveys to the model.

| Analysis group | Model / display name | Code name | Included features | No. features |
|---|---|---|---|---|
| Primary feature-block comparison | **ESM-2 only** | *llr_only* | 8 ESM-2 features | 8 |
| Primary feature-block comparison | **Site context only** | *evolution_site_only* | 4 site-level evolutionary features | 4 |
| Primary feature-block comparison | **Evolutionary context** | *evolution_only* | 4 site-level evolutionary features + 4 candidate-history features | 8 |
| Primary feature-block comparison | **Structure + functional annotations** | *structure_only* | 2 stability features + 5 binding features + 4 functional annotations | 11 |
| Primary feature-block comparison | **ESM-2 + evolutionary context** | *llr_evolution* | 8 ESM-2 features + 4 site-level evolutionary features + 4 candidate-history features | 16 |
| Primary feature-block comparison | **ESM-2 + structure / functional annotations** | *llr_structure* | 8 ESM-2 features + 2 stability features + 5 binding features + 4 functional annotations | 19 |
| Primary feature-block comparison | **Evolutionary context + structure / functional annotations** | *evolution_structure* | 4 site-level evolutionary features + 4 candidate-history features + 2 stability features + 5 binding features + 4 functional annotations | 19 |
| Primary feature-block comparison | **Molecular + site context** | *full_no_candidate_history* | 8 ESM-2 features + 4 site-level evolutionary features + 2 stability features + 5 binding features + 4 functional annotations | 23 |
| Primary feature-block comparison | **Complete model** | *full* | 8 ESM-2 features + 4 site-level evolutionary features + 4 candidate-history features + 2 stability features + 5 binding features + 4 functional annotations | 27 |
| Control | **Coverage-only control** | *coverage_only* | Indicators of ESM-2, stability and binding feature availability | 3 |
| DMS analysis | **Stability only** | *foldx_only* | 2 stability features | 2 |
| DMS analysis | **Binding only** | *rosetta_only* | 5 binding features | 5 |
| DMS analysis | **Stability + binding** | *foldx_rosetta* | 2 stability features + 5 binding features | 7 |
| DMS analysis | **ESM-2 + stability** | *llr_foldx* | 8 ESM-2 features + 2 stability features | 10 |
| DMS analysis | **ESM-2 + binding** | *llr_rosetta* | 8 ESM-2 features + 5 binding features | 13 |
| DMS analysis | **ESM-2 + stability + binding** | *llr_foldx_rosetta* | 8 ESM-2 features + 2 stability features + 5 binding features | 15 |
| DMS analysis | **Complete model without DMS** | *full_no_dms* | 8 ESM-2 features + 4 site-level evolutionary features + 4 candidate-history features + 4 functional annotations | 20 |
| DMS analysis | **Complete model without stability** | *full_no_foldx* | 8 ESM-2 features + 4 site-level evolutionary features + 4 candidate-history features + 5 binding features + 4 functional annotations | 25 |
| DMS analysis | **Complete model without binding** | *full_no_rosetta* | 8 ESM-2 features + 4 site-level evolutionary features + 4 candidate-history features + 2 stability features + 4 functional annotations | 22 |

**Table S2: Complete feature groups used in NoroScope model analysis.** Includes all groups of mathematical features used in training and validation runs of the NoroScope model. Each feature group is accompanied by its broad analysis group, name used in the NoroScope codebase, and a description of the number of features from each feature block included.

| Cutoff year | Feature combination | Average precision | ROC-AUC | Mean positive rank | Top-five recovery (%) |
|---|---|---|---|---|---|
| 2011 | **Complete model** | **0.805** | **0.856** | **3.02** | **87.9** |
| 2011 | **Evolutionary context** | 0.753 | 0.837 | 4.05 | 84.1 |
| 2011 | **ESM-2 + evolutionary context** | 0.783 | 0.845 | 3.17 | 83.3 |
| 2011 | **Molecular + site context** | 0.589 | 0.733 | 5.08 | 65.2 |
| 2011 | **ESM-2 + structure / functional annotations** | 0.558 | 0.706 | 5.16 | 65.2 |
| 2011 | **ESM-2 only** | 0.479 | 0.633 | 6.93 | 43.9 |
| 2011 | **Site context only** | 0.567 | 0.689 | 10.90 | 18.9 |
| 2011 | **Coverage-only control** | 0.402 | 0.523 | 10.90 | 18.9 |
| 2016 | **Complete model** | **0.834** | **0.904** | **2.31** | **92.9** |
| 2016 | **Evolutionary context** | 0.841 | 0.891 | 3.01 | 91.3 |
| 2016 | **ESM-2 + evolutionary context** | 0.839 | 0.901 | 2.26 | 93.4 |
| 2016 | **Molecular + site context** | 0.603 | 0.800 | 4.44 | 68.3 |
| 2016 | **ESM-2 + structure / functional annotations** | 0.529 | 0.748 | 4.93 | 64.5 |
| 2016 | **ESM-2 only** | 0.430 | 0.691 | 6.08 | 56.3 |
| 2016 | **Site context only** | 0.559 | 0.771 | 10.97 | 17.5 |
| 2016 | **Coverage-only control** | 0.346 | 0.564 | 10.97 | 17.5 |

**Table S3: Primary model performance across NoroScope feature combinations.** Performance was evaluated using pooled five-fold position-held-out out-of-fold predictions for the primary model cohort of historically-observed, non-dominant candidate amino acids. Average precision and ROC-AUC describe global classification, while mean positive rank and top-five recovery describe within-position prioritisation against all non-dominant amino acid alternatives. Positive prevalence was 37.6% in 2011 and 31.0% in 2016. For random within-position rankings, the expected mean rank is 10 and expected top-five recovery is 26.3%.

| Cutoff | Model / feature combination | Average precision | ROC-AUC | Mean positive rank | Top-five recovery (%) |
|---|---|---|---|---|---|
| 2011 | **Stability only** | 0.438 | 0.569 | 8.60 | 40.2 |
| 2011 | **Binding only** | 0.470 | 0.580 | 8.56 | 36.3 |
| 2011 | **Stability + binding** | 0.472 | 0.561 | 7.33 | 44.1 |
| 2011 | **ESM-2 only** | 0.545 | 0.619 | 7.58 | 43.1 |
| 2011 | **ESM-2 + stability** | 0.521 | 0.637 | 7.11 | 48.0 |
| 2011 | **ESM-2 + binding** | 0.520 | 0.617 | 6.91 | 50.0 |
| 2011 | **ESM-2 + stability + binding** | 0.495 | 0.621 | 6.49 | 52.0 |
| 2011 | **Complete model without DMS** | 0.757 | 0.796 | 4.14 | 77.5 |
| 2011 | **Complete model without stability** | 0.784 | 0.808 | 3.47 | 82.4 |
| 2011 | **Complete model without binding** | 0.777 | 0.807 | 3.94 | 81.4 |
| **2011** | **Complete model** | **0.786** | **0.811** | **3.36** | **82.4** |
| 2016 | **Stability only** | 0.450 | 0.651 | 8.08 | 39.3 |
| 2016 | **Binding only** | 0.478 | 0.554 | 9.52 | 34.3 |
| 2016 | **Stability + binding** | 0.490 | 0.658 | 7.54 | 46.4 |
| 2016 | **ESM-2 only** | 0.448 | 0.660 | 6.73 | 50.7 |
| 2016 | **ESM-2 + stability** | 0.481 | 0.701 | 6.05 | 57.1 |
| 2016 | **ESM-2 + binding** | 0.523 | 0.686 | 6.46 | 53.6 |
| 2016 | **ESM-2 + stability + binding** | 0.511 | 0.712 | 6.01 | 57.9 |
| 2016 | **Complete model without DMS** | 0.854 | 0.910 | 2.07 | 96.4 |
| 2016 | **Complete model without stability** | 0.863 | 0.918 | 2.12 | 97.1 |
| 2016 | **Complete model without binding** | 0.872 | 0.928 | 2.04 | 95.0 |
| **2016** | **Complete model** | **0.864** | **0.923** | **2.12** | **95.7** |

**Table S4: Ablation analysis for *in silico* DMS features.** Performance was evaluated using pooled five-fold position-held-out out-of-fold predictions using only positions in the primary cohort with structural DMS coverage, corresponding to the VP1 P domain. Average precision and ROC-AUC describe global classification, while mean positive rank and top-five recovery describe within-position prioritisation against all non-dominant amino acid alternatives. The 2011 analysis contained 247 modelling candidates across 137 positions (102 positives with 41.3% prevalence), and the 2016 analysis contained 396 candidates across 205 positions (140 positives with 35.4% prevalence). For random within-position rankings, expected mean rank is 10 and expected top-five recovery is 26.3%.

| Evaluation setting | Cutoff | Supported candidates | Positions with supported candidates | Mean positive rank | Top-five recovery (%) |
|---|---|---|---|---|---|
| Primary observed-alternative cohort | 2011 | 132 | 93 | 3.02 | 87.9 |
| Primary observed-alternative cohort | 2016 | 183 | 127 | 2.31 | 92.9 |
| Epitope positions | 2011 | 61 | 32 | 3.62 | 85.2 |
| Epitope positions | 2016 | 73 | 36 | 2.47 | 94.5 |
| HBGA-binding positions | 2011 | 20 | 10 | 4.65 | 75.0 |
| HBGA-binding positions | 2016 | 26 | 12 | 2.12 | 100.0 |
| Breathing-core positions | 2011 | 1 | 1 | 1.00 | 100.0 |
| Breathing-core positions | 2016 | 2 | 2 | 1.00 | 100.0 |
| **Historically unseen alternatives** | **2011** | **30** | **29** | **6.64 ± 3.05** | **52.7 ± 26.5** |
| **Historically unseen alternatives** | **2016** | **15** | **14** | **10.93 ± 2.76** | **23.3 ± 14.9** |
| Positions outside primary modelling cohort | 2011 | 13 | 13 | 9.23 | 46.2 |
| Positions outside primary modelling cohort | 2016 | 3 | 3 | 9.67 | 33.3 |

**Table S5: NoroScope performance across primary and secondary prediction tasks.** Performance was evaluated using pooled five-fold position-held-out out-of-fold predictions within both the primary modelling cohort and historical alternative prediction task as well as secondary analyses representing annotated functional regions and performance on *de novo* mutations. Annotated positions comprised epitopes A-I, the HBGA binding pocket and the breathing core as defined in table 1. The historically unseen (*de novo*) analysis was performed separately using candidates not observed before each time cutoff and five repeated position-grouped splits, hence values are reported as mean ± SD across model runs. For positions absent from the primary modelling cohort (due to a lack of historically observed alternative amino acids at the year cutoff), the primary cohort full model was used to score with results averaged across the five fitted models. No positions from outside the modelling cohort contributed to any of the fitted models. Positions which are position-annotated or contain historically unseen substitutions are not mutually exclusive with the primary model cohort. Random ranking among 19 alternative amino acids gives an expected mean rank of 10 and top-five recovery of 26.3%.

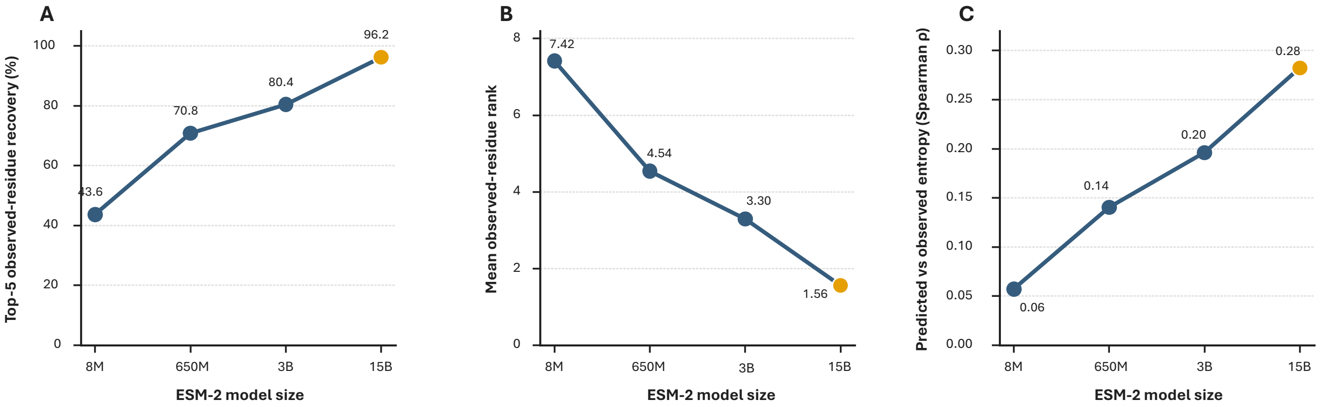


**Figure S1: ESM-2 model selection for zero-shot mutation analysis.** Performance of the 8M, 650M, 3B and 15B pre-trained ESM-2 models was evaluated across 11 representative VP1 sequence contexts. Models were compared using **A)** top-five recovery of observed alternative amino acids, **B)** mean within-position rank of observed alternative amino acids, and **C)** the Spearman correlation between ESM-2 site entropy and the observed sequence entropy. In all cases, ESM-2 15B achieved the highest performance, with overall performance increasing with model size.

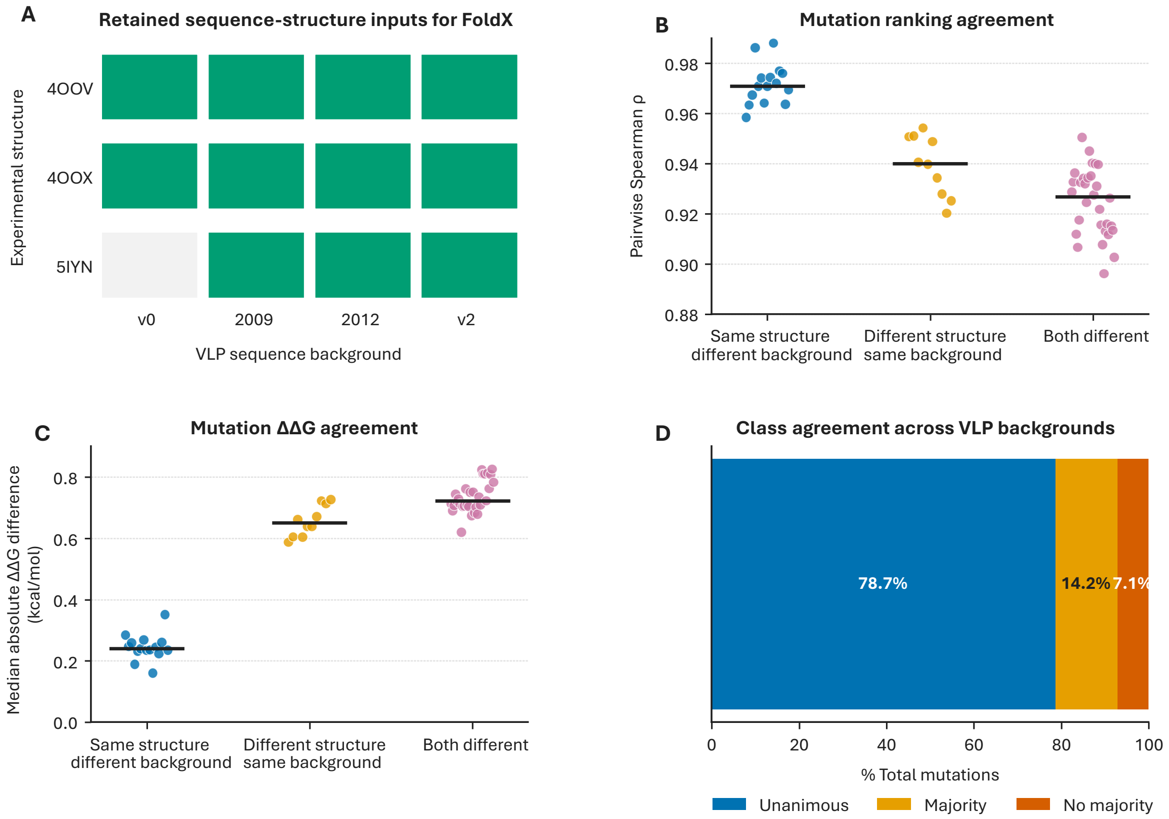


**Figure S2: Technical quality control of the FoldX-derived protein stability deep mutational scanning dataset. A)** Combinations of experimental structures mutated to match VLP sequences used in the NoroScope database. 5IYN mutated to match v0 was the only excluded structure-sequence combination due to being a technical outlier. **B)** Pairwise Spearman rank correlations of mutation-level FoldX ΔΔG values between included inputs, grouped by whether comparisons share the same experimental structure, the same VLP sequence background, or differ in both. **C)** The median absolute difference in ΔΔG for mutations across the same comparison groups, illustrating the difference in predicted effect magnitude between structural contexts. **D)** Stability-effect class agreement across available VLP sequence backgrounds, summarised as unanimous agreement, majority agreement, or no majority.

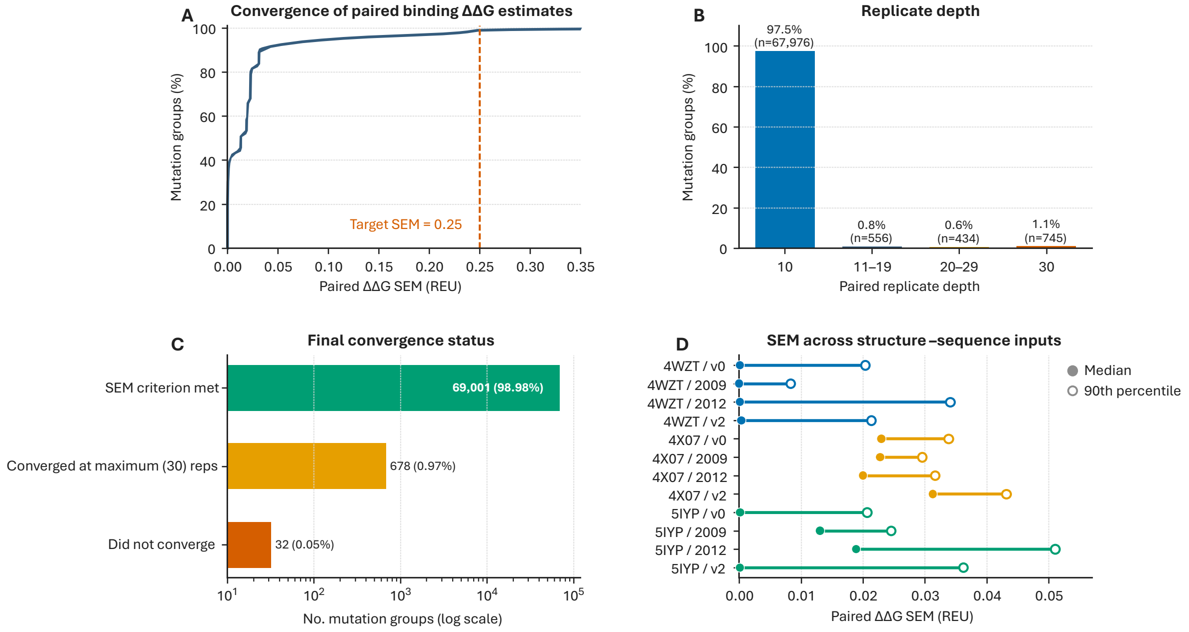


**Figure S3: Technical quality control of the Rosetta-derived HBGA binding deep mutational scanning dataset. A)** Cumulative distribution of the SEM of paired $\Delta\Delta G_{bind}$ estimates across all mutation-template calculations, with the defined converge threshold of 0.25 REU indicated by the dashed line. **B)** Distribution of paired mutations after adaptive repeats, showing that the majority of mutation-template combinations converged to the target SEM of 0.25 REU within 10 replicates. **C)** Final convergence status of all 69,711 mutation-template combinations. 98.98% of mutations converged without reaching the replicate limit, while 0.97% converged while hitting the limit and 0.05% did not converge. **D)** Median and 90$^{th}$ percentile SEM values for each structure-sequence input used in the binding analysis, demonstrating consistently low uncertainty across all 12 combinations.

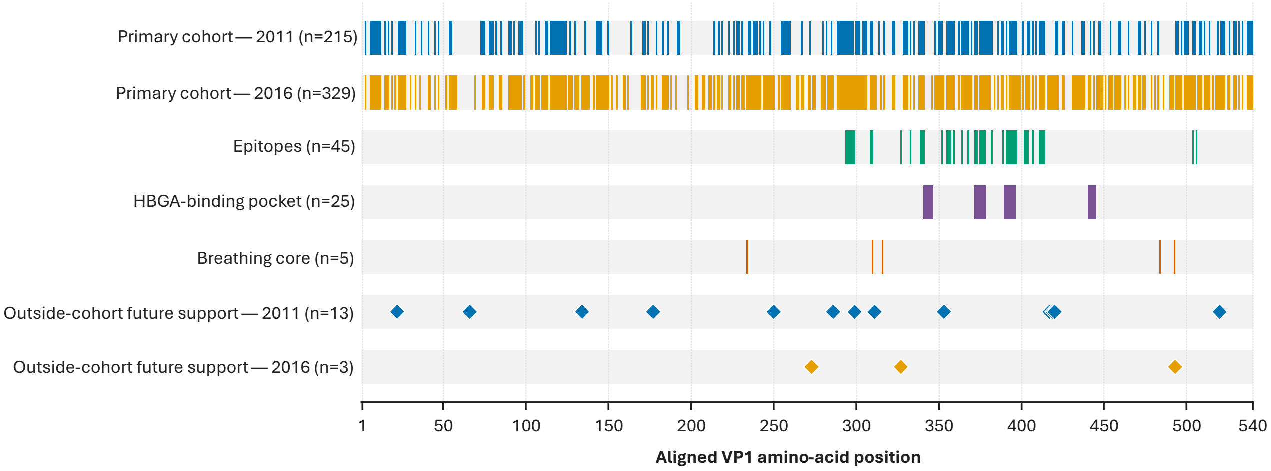


**Figure S4: Positional coverage of the primary NoroScope modelling cohort across HuNoV GII.4 VP1.** A position was included within the primary modelling cohort if it contained at least one historically observed alternative amino acid to the most frequent (dominant) residue. Positions included within the primary modelling cohorts for the 2011 and 2016 cutoffs are displayed across the aligned VP1 sequence used in the NoroScope database. Positions not included within the primary modelling cohort at which at least one future-supported mutation subsequently arose are displayed separately. Positions corresponding to annotated functional regions (epitopes, the HBGA binding pocket and the breathing core) are also displayed.
